# An annotated biobank of triple negative breast cancer patient-derived xenografts featuring treatment-naïve and longitudinal samples throughout neoadjuvant chemotherapy

**DOI:** 10.1101/2024.11.25.625287

**Authors:** Amanda L. Rinkenbaugh, Yuan Qi, Shirong Cai, Jiansu Shao, Faiza Hancock, Sabrina Jeter-Jones, Xiaomei Zhang, Emily Powell, Lei Huo, Rosanna Lau, Chunxiao Fu, Rebekah Gould, Petra den Hollander, Elizabeth E. Ravenberg, Jason B. White, Gaiane M. Rauch, Banu Arun, Clinton Yam, Alastair M. Thompson, Gloria V. Echeverria, Stacy L. Moulder, W. Fraser Symmans, Jeffrey T. Chang, Helen Piwnica-Worms

## Abstract

Triple negative breast cancer (TNBC) that fails to respond to neoadjuvant chemotherapy (NACT) can be lethal. Developing effective strategies to eradicate chemoresistant disease requires experimental models that recapitulate the heterogeneity characteristic of TNBC. To that end, we established a biobank of 92 orthotopic patient-derived xenograft (PDX) models of TNBC from the tumors of 75 patients enrolled in the ARTEMIS clinical trial (NCT02276443) at MD Anderson Cancer Center, including 12 longitudinal sets generated from serial patient biopsies collected throughout NACT and from metastatic disease. Models were established from both chemosensitive and chemoresistant tumors, and nearly 30% of PDX models were capable of lung metastasis. Comprehensive molecular profiling demonstrated conservation of genomes and transcriptomes between patient and corresponding PDX tumors, with representation of all major transcriptional subtypes. Transcriptional changes observed in the longitudinal PDX models highlight dysregulation in pathways associated with DNA integrity, extracellular matrix interactions, the ubiquitin-proteasome system, epigenetics, and inflammatory signaling. These alterations reveal a complex network of adaptations associated with chemoresistance. This PDX biobank provides a valuable resource for tackling the most pressing issues facing the clinical management of TNBC, namely chemoresistance and metastasis.

## INTRODUCTION

Triple negative breast cancer (TNBC) is a subtype of breast cancer characterized by the absence of estrogen and progesterone hormone receptor expression and the lack of amplification or overexpression of *ERBB2*, encoding human epidermal growth factor receptor 2. The lack of these receptors precludes the use of some of the most effective targeted breast cancer treatments for these patients. Currently, systemic neoadjuvant chemotherapy (NACT) is the established treatment approach for early stage TNBC^1^, with recent approval for immunotherapy in the neoadjuvant setting^2^. However, up 40-50% of patients with TNBC are found to have residual disease post-NACT, even with the addition of immunotherapy, leading to a heightened risk of early recurrence after surgical resection^1,3^. Identifying upfront predictors of response to NACT remains a critical clinical challenge, as it could help tailor treatments to those that would be most effective with the least toxicity.

To understand response to NACT, the ARTEMIS trial (A Robust TNBC Evaluation FraMework to Improve Survival, NCT02276443) evaluated whether a gene expression signature^4^ could be used to predict patients with chemoresistant disease and direct them to phase II clinical trials of targeted therapies based on biomarkers from their pre-treatment tumors. Tumor tissue was collected at multiple clinical timepoints: before NACT initiation, from tumors showing a poor response after four cycles of doxorubicin and cyclophosphamide (AC), at surgical resection following completion of NACT, and from metastatic disease. Patient tumors were profiled by whole exome sequencing (WES) and transcriptome sequencing (RNA-seq). It is important to note that the ARTEMIS trial preceded KEYNOTE-522^2,5,6^, and therefore none of the patients received immunotherapy as part of the standard of care treatment, although some patients were enrolled in a phase II trial combining atezolizumab with nab-paclitaxel^7^.

Patient-derived xenograft (PDX) models have been established as generally faithful recapitulations of their source tumors, providing a platform for both experimental and translational research^8–13^. To support translational research in TNBC and mitigate the challenges posed by limited patient tumor tissue, we established a biobank of orthotopic PDX models from tumors collected in the ARTEMIS trial. The biobank includes 92 PDX models derived from the tumors of 75 patients, including 12 longitudinal sets established from serially sampled tumors across trial timepoints or metastatic lesions. The models encompassed a range of TNBC subtypes. A high degree of conservation in genomes, transcriptomes, and molecular subtypes was observed between PDX tumors and their respective patient tumors based on WES and RNA-Seq. The genomic profiles of the PDX tumors accurately reflected the landscape of TNBC, with *TP53* being the most frequently observed genetic alteration. In addition, lung metastases were detected in a significant fraction of PDX models. Thus, this annotated PDX biobank comprehensively captures the genomic and transcriptional diversity of TNBC, including representation of each molecular subtype, and promises to be a valuable resource for addressing critical challenges facing TNBC patients with chemoresistant and metastatic disease.

## RESULTS

### Generation of a TNBC PDX biobank in alignment with the ARTEMIS trial

Patients with TNBC enrolled in the prospective neoadjuvant ARTEMIS trial (NCT02276443) provided tumor material for WES, RNA-seq, histology, and establishment of PDX models via implantation into mice. Longitudinal patient samples were collected at different stages of neoadjuvant treatment: prior to treatment (pre-NACT), after four cycles of doxorubicin and cyclophosphamide (AC; mid-NACT), following treatment with paclitaxel, alone or in combination with an experimental targeted therapy (post-NACT), and from metastatic disease (Figure 1A). For each patient sample, we pooled cells from three independent fine needle aspirations (FNAs) and engrafted them into the pre-humanized fourth mammary fat pads (MFPs) of non-obese diabetic/severe combined immunodeficient (NOD/SCID) mice as described previously^8,14,15^. We classified PDX models as established once they reached the third passage and consistently passed quality control assays. These tests included short-tandem repeat (STR) DNA fingerprinting to verify the unique tumor profile and corresponding patient match of each tumor, quantitative PCR (qPCR) using human and mouse-specific DNA probes to confirm the human origin of tumor cells, and PCR to confirm the absence of GFP, which was present in immortalized human stromal fibroblasts used to humanize the mouse MFPs before the initial engraftment of patient tumor cells.

**Figure 1:**
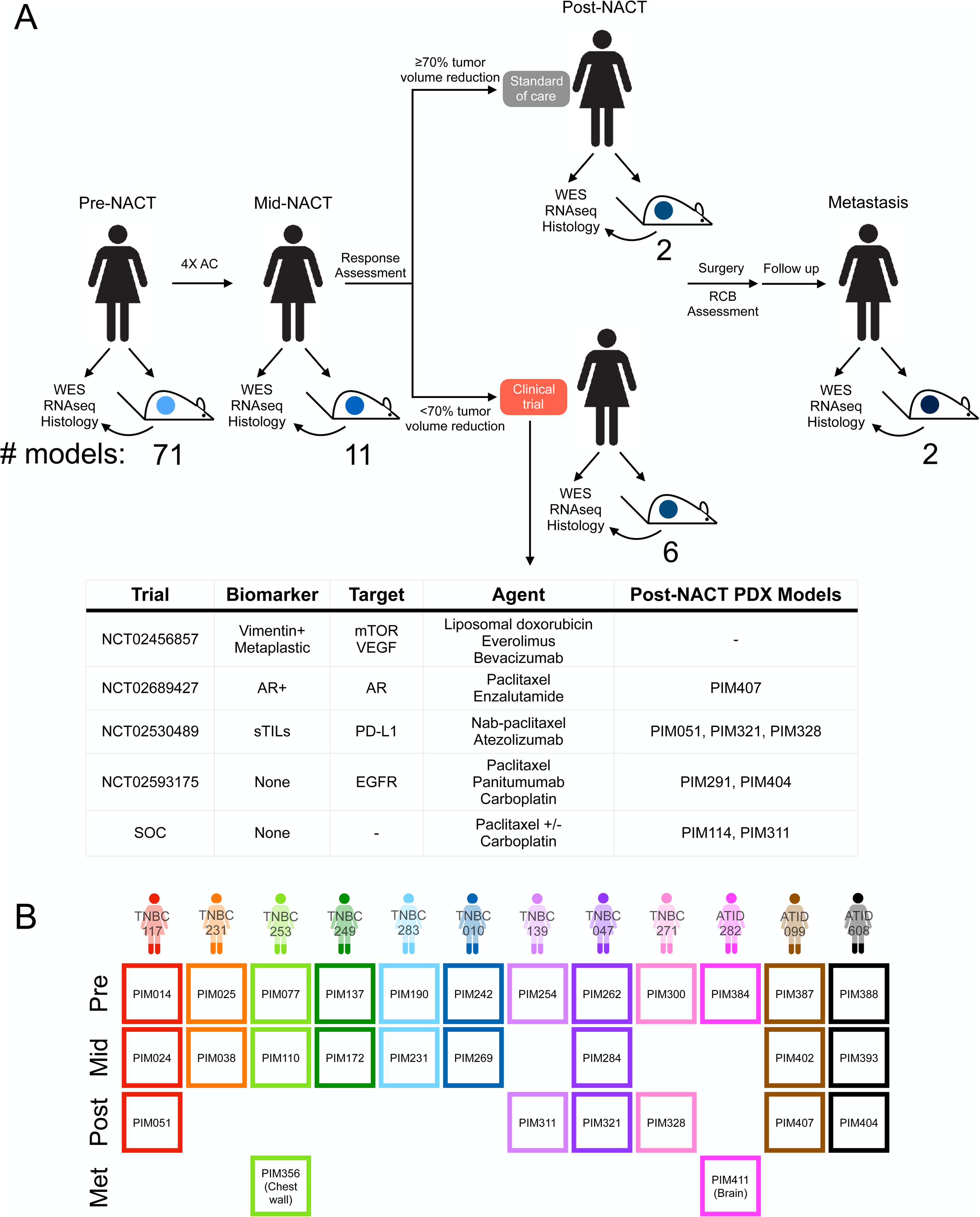
Generation of a TNBC PDX biobank in alignment with the ARTEMIS trial. (A) Schematic of ARTEMIS trial design where all treatment-naïve patients received four cycles of AC, followed by a response assessment via ultrasound or MRI. Patients with ≥70% tumor volume reduction remained on standard of care treatment, while patients with <70% tumor volume reduction were offered one of four clinical trials. The included table describes the associated phase II clinical trials that offered targeted therapies to patients with suboptimal AC response. Samples were collected from patients at key timepoints (pre-, mid-, and post-NACT and metastasis) to be analyzed by WES, RNA-seq, and histology, as well as implanted into mice for PDX model generation. (B) Visualization of 12 longitudinal sets encompassing pre-, mid-, and post-NACT, and metastatic samples. Patient identifier is shown above each column of corresponding samples. Biopsy timepoint is indicated to the left of each row. AC: doxorubicin and cyclophosphamide, RCB: residual cancer burden, WES: whole exome sequencing,

Out of the 92 established PDX models derived from tumors of 75 patients, 71 originated from pre-NACT biopsies, 11 from mid-NACT biopsies, eight from post-NACT biopsies and two from metastases (PIM356: chest wall, PIM411: brain). Additionally, the collection includes 12 longitudinal sets, each featuring a pre-NACT model along with at least one additional model from mid-NACT, post-NACT, or metastasis (Figure 1B). As part of ARTEMIS, four phase II therapeutic trials (Figure 1A) were available to explore targeted therapies in combination with paclitaxel for those patients with suboptimal responses to AC, based on tumor volume reduction by ultrasound or MRI and clinical assessment. Eligibility for the phase II trials followed a hierarchical selection process based on specific tumor characteristics determined at diagnosis. Patients with tumors exhibiting ≥20% tumor infiltrating lymphocytes (TILs) were eligible for treatment with atezolizumab and nab-paclitaxel (NCT02530489)^7^, then tumors with metaplastic histology or vimentin IHC expression >50% were eligible for liposomal doxorubicin and bevacizumab combined with temsirolimus or everolimus (NCT02456857)^16^, then tumors with androgen receptor IHC expression ≥10% were eligible for enzalutamide and paclitaxel (NCT02689427)^17^. If none of these biomarkers were present, patients were eligible for panitumumab in combination with carboplatin and paclitaxel (NCT02593175)^18^. Of the eight PDX models established from post-NACT tumors, two were from patients who received paclitaxel as standard of care (PIM114, PIM311); three were from patients enrolled in NCT02530489 (PIM051, PIM321, PIM328); one was from a patient enrolled in NCT02689427 (PIM407); and two were from patients enrolled in NCT02593175 (PIM291 and PIM404) (Figure 1A). The PDX collection was established from a diverse patient population with 21% identifying as black, 16% as Hispanic or Latino, and 7% as Asian (Table S1). Histological analysis demonstrated that 96% of the PDX tumors were invasive ductal carcinoma, while the remaining four models were classified as metaplastic tumors. Longitudinal PDX models generally retained the same histology across timepoints and shared the histology of the corresponding patient samples (Figure S1)^19^. Detailed clinical annotation and characterization of each PDX model can be found in Table S1.

We collected 392 patient tumors to engraft into mice, which produced the final collection of 92 models resulting in a take rate of 23%, in agreement with our previously reported interim analysis^14^. In that study, we also found that therapy-resistant tumors, defined either as RCB-II/III after NACT or with metastatic relapse within two years of surgery, were positively correlated with successful PDX engraftment^14^. Assessing the entire biobank, we confirmed that both RCB-II/RCB-III tumors and tumors with metastatic relapse more frequently led to successful establishment of a PDX model. The recent prospective clinical trial TOWARDS further underscored the relationship between successful PDX engraftment and metastatic relapse^20^. The patient biopsy timepoint did not correlate with take rate (Figure S2A-2SF). Finally, we investigated the time required for each model to reach passage three and observed that higher RCB status correlated with a faster time to PDX establishment, independent of molecular subtype (Figure S2G-S2I).

Given that TNBC is known for its aggressive metastatic behavior, we investigated our PDX models for the presence of lung metastases at endpoints following primary tumor resection or declining health of the mice. We analyzed hematoxylin and eosin-stained lung sections, and in some cases, stained for cytokeratins and human mitochondria (Figure 2A). Our findings revealed that 28% of the examined models (21/75) could colonize the lungs of engrafted mice. Metastasis rates between transcriptional subtypes did not vary significantly perhaps due to sample size limitations, though BL1 and LAR trended towards higher rates (Figure 2B). There was no significant association between RCB status and the propensity for lung metastasis (Figure 2C). It is possible that we would have detected metastases in additional models if primary tumors were given more time *in vivo* or if more sensitive detection methods were employed. Based on rapid and overt lung metastases detected in PIM056, a pre-treatment model, we engineered it to express click beetle red-luciferase and used bioluminescence imaging to assess metastasis *in vivo.* We identified metastatic lesions in the lung, liver, brain, and bone through this approach (Figure 2D). These results agreed in part with the medical history of the patient who exhibited both lung and liver metastases. Multi-organ metastasis has previously been reported in other PDX models^11,21,22^, and it will be important to assess the extent to which additional PDX models in our collection are capable of multi-organ metastasis.

**Figure 2:**
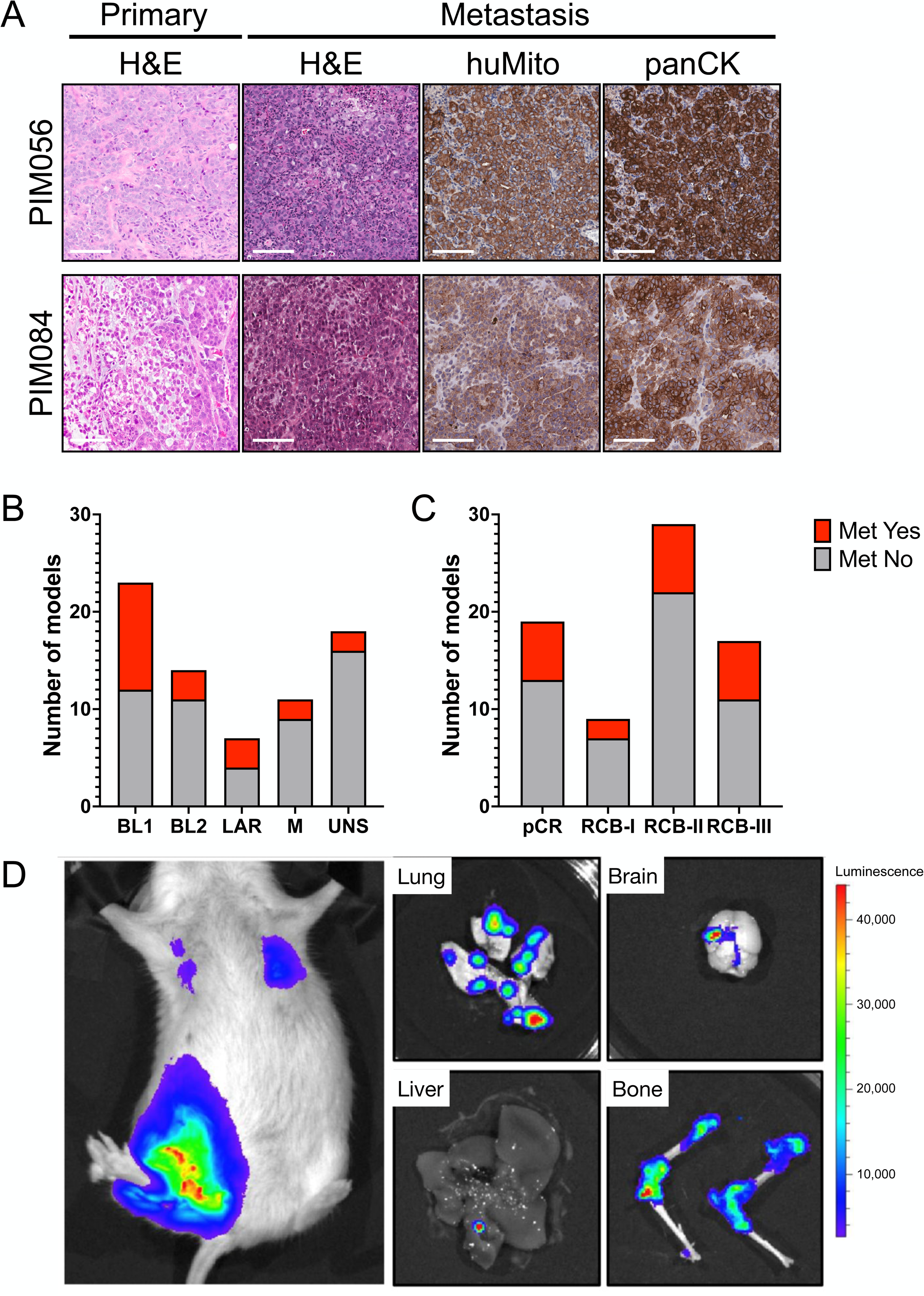
Lung metastasis is a common feature of TNBC PDX models. (A) H&E and IHC staining of pan-cytokeratin and human-specific mitochondria in the primary tumors and lung metastases from PIM056 and PIM084. Scale bars are 100 µm. (B) Quantification of metastasis-competent PDX models based on transcriptional subtype (C) Quantification of metastasis-competent PDX models based on RCB status (D) Bioluminescence imaging of PIM056 labeled with CBR-luciferase identifies metastases in the lung, liver, brain, and bone.

### Genomic and transcriptomic profiles of PDX tumors reflect the diversity of human TNBC

We profiled all 92 PDX models, representing the entire PDX collection, by WES and RNA-seq. *TP53* was the most frequently altered gene, with 50 tumors exhibiting a somatic mutation, including 70% missense, 22% stopgain, and 10% splicing mutations (Figure 3A), in line with other studies of TNBC^23–25^. To evaluate the distribution of somatic mutations in our PDX dataset, we compared mutation frequencies with those from the METABRIC TNBC cohort^26,27^, focusing on the top 20 most frequently mutated genes in that dataset. Fisher’s exact tests showed no significant difference in mutation frequencies between our PDX dataset and the METABRIC TNBC cohort, except for a lower prevalence of *TP53* mutations in our PDX models (Figure 3B), and their corresponding patient tumors.

**Figure 3:**
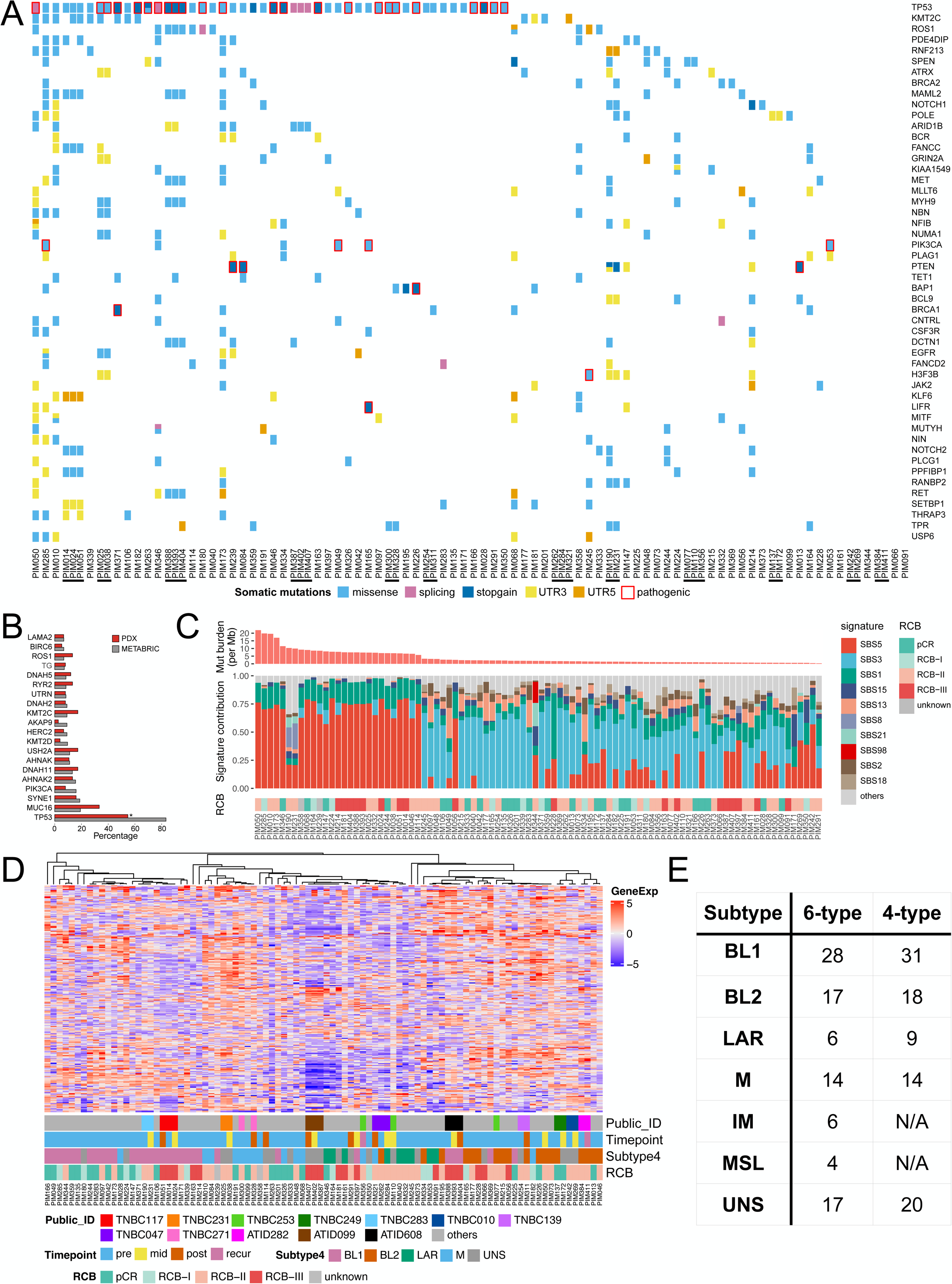
Genomic and transcriptomic profiles of PDX tumors reflect the diversity of human TNBC. (A) Oncoprint showing mutations of commonly mutated cancer genes from the COSMIC database. Models derived from the same patient are indicated by bars below the PDX ID. (B) Comparison of mutation frequency of the top 20 most frequently mutated genes in METABRIC TNBC samples and TNBC PDX biobank. (C) Stacked barplot shows the relative contributions of mutational signatures in each PDX model. Models are in order from highest to lowest overall mutation burden, reflected in bar graph at the top. RCB status is annotated at the bottom. (D) Heatmap of the 250 most variable genes across the RNAseq dataset of all 92 PDX models. Annotations for each PDX model include patient ID, transcriptional subtype, RCB status, and biopsy timepoint. (E) Summary of the number of PDX models assigned to each transcriptional subtype for a 6-class or 4-class system

We also found multiple pathogenic alterations in the PI3K pathway, including four *PIK3CA* missense mutations (E542K in PIM049, PIM165, and PIM285; H1047R in PIM053) and three nonsense *PTEN* mutations. None of the patients had germline mutations in *BRCA1/2* as those patients were recruited to a different clinical trial at MD Anderson Cancer Center^28,29^. However, five models had somatic *BRCA1* mutations, and nine models had somatic *BRCA2* mutations. While most of these mutations were classified as benign or variants of unknown significance, PIM371 harbored a stopgain *BRCA1* mutation (R1443X) that is reported to be deleterious^30^. Comparing the longitudinal models, we found that 96 out of 213 non-silent mutations in COSMIC genes were consistently present across all timepoints. Notably, PIM190 and PIM231, established from the pre- and mid-NACT samples of patient TNBC283, exhibited the greatest discordance. This discrepancy aligns with the variability observed in STR profiling across multiple passages of these models, where the number of microsatellite repeats fluctuated within the acceptable range of the assay, allowing both models to still be assigned the same profile. We assigned the PDX models into one of three categories based on their tumor mutation burden: high (>15 non-synonymous somatic mutations/Mb), medium (5-15 mutations/Mb), and low (<5 mutations/Mb). 71% of tumors fell in the low category, 25% in the medium category, and 4% in the high category (Figure 3C, top). Analysis of the mutational signatures showed that the high and medium mutation burden tumors were dominated by signature SBS5, while the low mutation burden tumors were characterized by signature SBS3 (Figure 3C, bottom)^31^. Signature SBS5 is associated with a clock-like etiology, while signature SBS3 is linked with defects in homologous recombination repair. Neither signature aligned with readily discernible characteristics of the patients (such as age) or their corresponding PDX models (such as BRCA status).

Additionally, we used WES data to generate genome-wide copy number profiles for each PDX model. We observed frequent gains of chromosomes 1q and 8q, as well as losses on chromosome 5, which is consistent with previous findings in human TNBC (Figure S3). Comparing longitudinal changes across the matched sets, we noted a gain of 16p in five of the mid- or post-NACT models. The gain of 16p has been linked to increased immune evasion^32^, which may indicate an adaptive response of these tumors to treatment.

Based on transcriptional profiling, we clustered the PDX models using the top 250 most variable genes. We found that 11 out of 12 longitudinal sets were closely clustered together, suggesting that patient-level heterogeneity is a more significant factor influencing transcriptional state than treatment timepoint (Figure 3D). To assign transcriptional subtypes to each tumor, we utilized TNBCtype, opting for the 4-type^33^ over the 6-type^34^ classifier. The 4-type classifier better reflects cancer cell properties and minimizes stromal influences, which is better aligned with our gene expression profiles since we discarded the host stromal reads when processing the data from PDX tumors. All four molecular subtypes of TNBC, basal-like 1 (BL1), basal-like 2 (BL2), luminal androgen receptor (LAR), and mesenchymal (M)^33,34^, are represented in our PDX collection (Figure 3E). Additionally, our analysis revealed significant intratumoral heterogeneity when examining subtype coefficients as continuous variables. Specifically, 34 out of 92 (37%) of the PDX models scored significantly for more than one subtype (Figures S4, S7). For tumors with multiple significant subtype coefficients, we found that the BL1 and M subtypes significantly co-occurred (p = 0.005), while the BL2 and LAR subtypes were the next most frequently co-occurring but did not reach statistical significance (p = 0.13).

### PDX models conserve genotype and phenotype of patient samples

After analyzing the genomic and transcriptional landscape of our PDX biobank, we next assessed how well the PDX models preserved the features of the corresponding patient tumors. WES data was available for 71 out of 75 patients, accounting for 87 PDX tumors and 11 of the 12 longitudinal sets, while RNA-seq data was available from 70 patients, accounting for 84 PDX tumors and 10 of 12 longitudinal sets (Table S2). Our analysis of conserved mutations between patient tumor and corresponding PDX tumor revealed a median conservation rate of 0.55, and no correlation with tumors from non-corresponding patients (Figure 4A). As *TP53* was the most frequently mutated gene, we specifically examined if *TP53* mutations were conserved between patient and PDX tumors. There was a 94% concordance in *TP53* alterations between patient and PDX tumors. Exceptions included PIM166 and PIM263 with pathogenic Y220C and G245D mutations respectively, not found in the corresponding patient tumors, and PIM056, which lacked the pathogenic R342X mutation detected in the patient tumor. Additionally, the P72R mutation present in the pre-NACT patient tumor from TNBC249 was not detected in the mid-NACT patient tumor or the associated PDX models, PIM137 (pre-NACT) or PIM172 (mid-NACT). P72R is a common *TP53* polymorphism, which is characterized as benign in ClinVar but has been associated with increased metastasis and poorer prognosis in p53-mutant breast cancer^35,36^.

**Figure 4:**
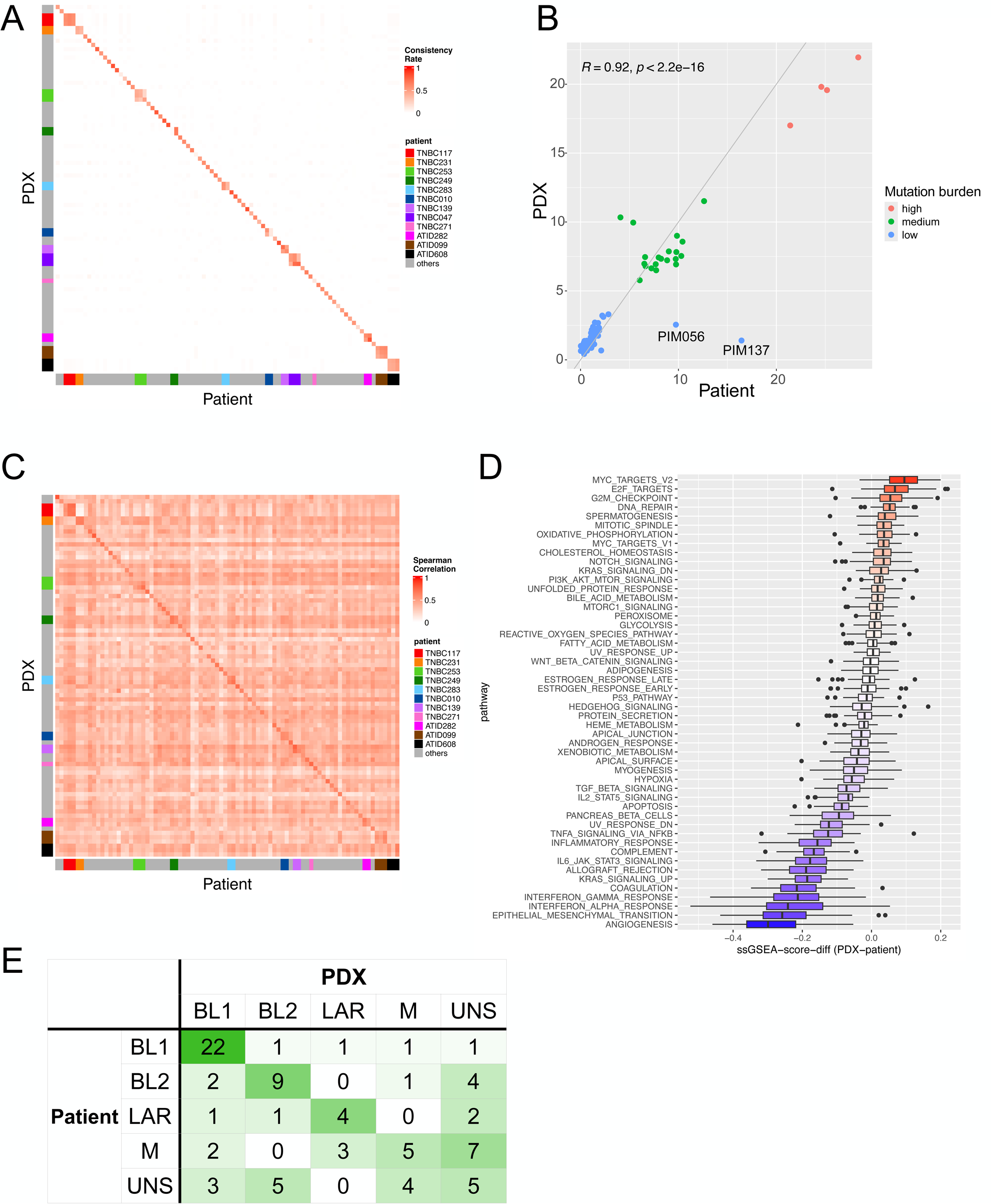
PDX models conserve genotype and phenotype of patient samples. (A) Heatmap of consistency rates for mutations between patient samples and PDX models. Longitudinal sets of samples are indicated with patient color scheme. (B) Scatter plot showing the correlation between mutation burden in patient and PDX samples, with color scheme representing high, medium, and low mutation burden categories. (C) Heatmap of the Spearman correlation coefficients for the 1000 most variable genes between patient samples and PDX models. Longitudinal sets of samples are indicated with patient color scheme. (D) Differential ssGSEA pathway scores for the Hallmark pathways between patient samples and PDX models (E) Confusion matrix of subtype assignments between patient samples and PDX models.

We also assessed the overall mutation burden and discovered that 90 out of 92 PDX models (98%) were classified into the same category of mutation burden as their corresponding patient tumor (Figure 4B). The two exceptions, PIM056 and PIM137, may be attributed to variations in the subclonal populations sampled for sequencing and PDX model establishment, or the possibility that certain subclones, while detected in patient tumors, were unable to proliferate in mice.

Transcriptomic analysis via bulk RNA sequencing showed a median Spearman correlation of 0.59 for the 1000 most variable genes between patient and PDX tumors, indicating a notable preservation of expression patterns between patient tumors and their corresponding PDX tumors (Figure 4C). Gene set enrichment analysis (GSEA) of the 50 Hallmarks pathways^37^ revealed distinct patterns of pathway expression: patient samples exhibited higher expression of angiogenesis and inflammation-related pathways, whereas PDX tumors showed increased proliferation and DNA repair pathways (Figure 4D). These differences may be due, in part, to the inclusion of stromal transcriptomes in the patient samples that were removed from the PDX samples by discarding transcripts mapped to a mouse reference genome. Additionally, the predominance of the BL1 molecular subtype in the PDX collection (Figure 3E) likely contributes to the observed enrichment in proliferation and DNA repair pathways noted in the PDX samples. We also assessed conservation of transcriptional subtypes between patient and PDX tumors using the 4-type classification system. 54% of the most significant subtypes in the patient tumors were also the most significant in the corresponding PDX models, with the highest concordance in BL1 tumors (Figure 4E). As previously noted, the subtype assignments can be ambiguous, where tumors can obtain statistically significant scores for multiple subtypes. When comparing all statistically significant subtype assignments, we found a 71% concordance between patient and PDX tumor subtypes (Table S1).

While the PDX models generally recapitulated properties of their corresponding patient samples, some discrepancies were noted. To quantify their overall similarity, we calculated conservation scores across five categories: mutations, mutation burden, subclonal architecture, gene expression, and transcriptional subtype. Each PDX model was scored as 0 (weak conservation), 0.5 (moderate conservation), or 1 (strong conservation) in each category, with the total conservation score derived from the sum across all categories. There was no significant correlation between conservation scores and various biological variables, including treatment timepoint, mutation burden, RCB status, patient metastasis, or patient or PDX transcriptional subtype (Supplemental Figure 5A-F). Among the longitudinal sets, we found that seven out of nine PDX sets either maintained or improved their conservation score across treatment timepoints (Supplemental Figure 5G).

### Tumors within longitudinal PDX sets exhibit AC resistance

We conducted preclinical studies to evaluate the response of several longitudinal PDX tumor sets to AC (Figure 5). These sets included four pre-/mid-NACT pairs, two pre-/post-NACT pairs, and two pre-/mid-/post-NACT triplets. Mice engrafted with these tumors were treated with either vehicle or a single dose of AC at our previously determined maximum tolerated dose for NOD/SCID mice^38^, and tumor volumes were measured twice weekly for up to 90 days. As anticipated, all tumors exhibited resistance to AC, given that seven out of the eight sets were derived from resistant patient tumors (RCB-II/RCB-III). Although the patient whose tumor was used to establish the PIM137/PIM172 pair achieved a pCR on the phase II immunotherapy trial (NCT02530489)^7^, her tumor volume increased during AC treatment as measured by ultrasound, predicting it was AC-resistant.

**Figure 5:**
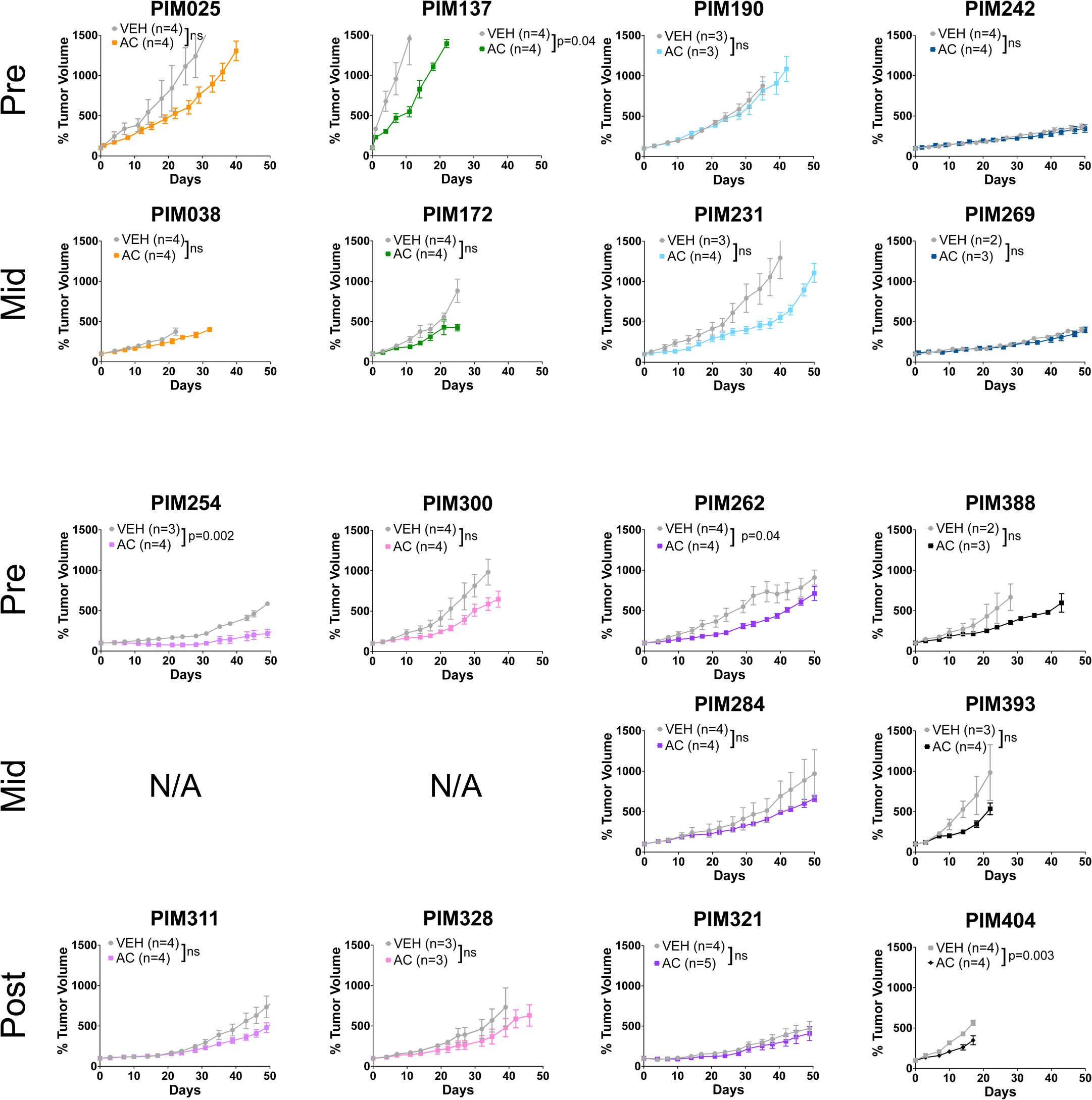
Tumors within longitudinal PDX sets exhibit AC resistance. Tumor response to one dose of AC on Day 0, showing mean tumor volume relative to starting tumor volume. Longitudinal sets are indicated by color scheme and model timepoint in indicated to the left of each row. Growth curves are based on n=2-5 mice/group, as indicated in the graph legends. Data are presented as the mean ± SEM. Statistical significance is reported based on unpaired two-tailed t-tests comparing the area under the curve values for vehicle vs. AC-treated tumors from treatment start until vehicle tumors reached endpoint.

The increase in tumor volume observed in most AC-treated mice was either similar or slightly slowed relative to the vehicle-treated controls (Figure 5), suggesting intrinsic resistance. In one case, PIM254, a pre-NACT tumor, initially regressed during AC treatment, but then ceased responding and resumed growth (Figure 5, bottom left). In contrast its corresponding post-NACT tumor, PIM311, grew continuously on AC treatment. This finding aligns with the patient’s treatment history, as the tumor showed a 95% decrease in volume at the mid-treatment timepoint but was classified as RCB-II at the time of surgery.

### Analysis of the subclonal architectures, transcriptomes, and proteomes of longitudinal PDX tumor sets

We analyzed both patient and PDX tumors before and during treatment to determine how chemotherapy impacted subclonal architecture. Using PyClone^39^, we estimated cancer cell frequency after accounting for tumor purity and local copy number status. This analysis did not reveal subclone enrichment after AC treatment in any of the patient or PDX tumors when comparing pre-NACT tumors to mid-NACT tumors (Figure S6). By contrast, the analysis of one patient tumor (TNBC139) showed subclone expansion in both the post-NACT tumor and its corresponding PDX tumor, PIM311 (Figure S6, bottom left). Notably, these samples correspond to the PIM254/PIM311 PDX pair with distinct AC responses. Additionally, two longitudinal sets included metastatic lesions: one (PIM411) paired with a pre-NACT tumor (PIM384), and the other (PIM356) paired with both pre- (PIM077) and mid-NACT (PIM110) tumors. In both cases, the metastatic lesions exhibited subclone selection relative to the earlier timepoints, consistent in both patient and PDX samples.

We analyzed the transcriptomes of tumors within each longitudinal PDX set to identify therapy-induced changes in the Hallmark pathways^37^. Table S3 summarizes data available for each model among the longitudinal sets. Each set of models exhibited a distinct set of altered pathways, reflecting the substantial patient-level heterogeneity seen in TNBC datasets^40,41^. In the mid-NACT models, the most commonly upregulated pathways relative to pre-NACT models could be grouped into three major categories: proliferation (E2F targets, MYC targets V1, G2M checkpoint), inflammatory signaling (interferon alpha and gamma responses, and inflammatory response), and epithelial-mesenchymal transition (Figure 6A). These pathways were upregulated in an overlapping, but not identical, subset of models with upregulation observed in 4-5 mid-/pre-NACT comparisons. Conversely, the most commonly downregulated pathways in mid-NACT compared to pre-NACT tumors were hypoxia and TNFɑ signaling via NF-κB. The dichotomy between upregulated interferon signaling and downregulated TNFɑ signaling may reflect two distinct types of chemotherapy response, as these signatures typically change in concert. Notably, only one mid-/pre-NACT comparison (TNBC231) showed these pathways significantly altered in opposite directions; in all other comparisons, either only one pathway was significantly changed, or both pathways moved in the same direction. To gain deeper insights into transcriptional changes over time, we also analyzed the samples using the MSigDB C2 pathways^42^. We identified 81 pathways that were significantly altered in at least 5 out of 9 instances, which fell into four main categories: extracellular matrix (ECM), ubiquitin-proteasome system, DNA integrity, and epigenetics (Figure 6B).

**Figure 6:**
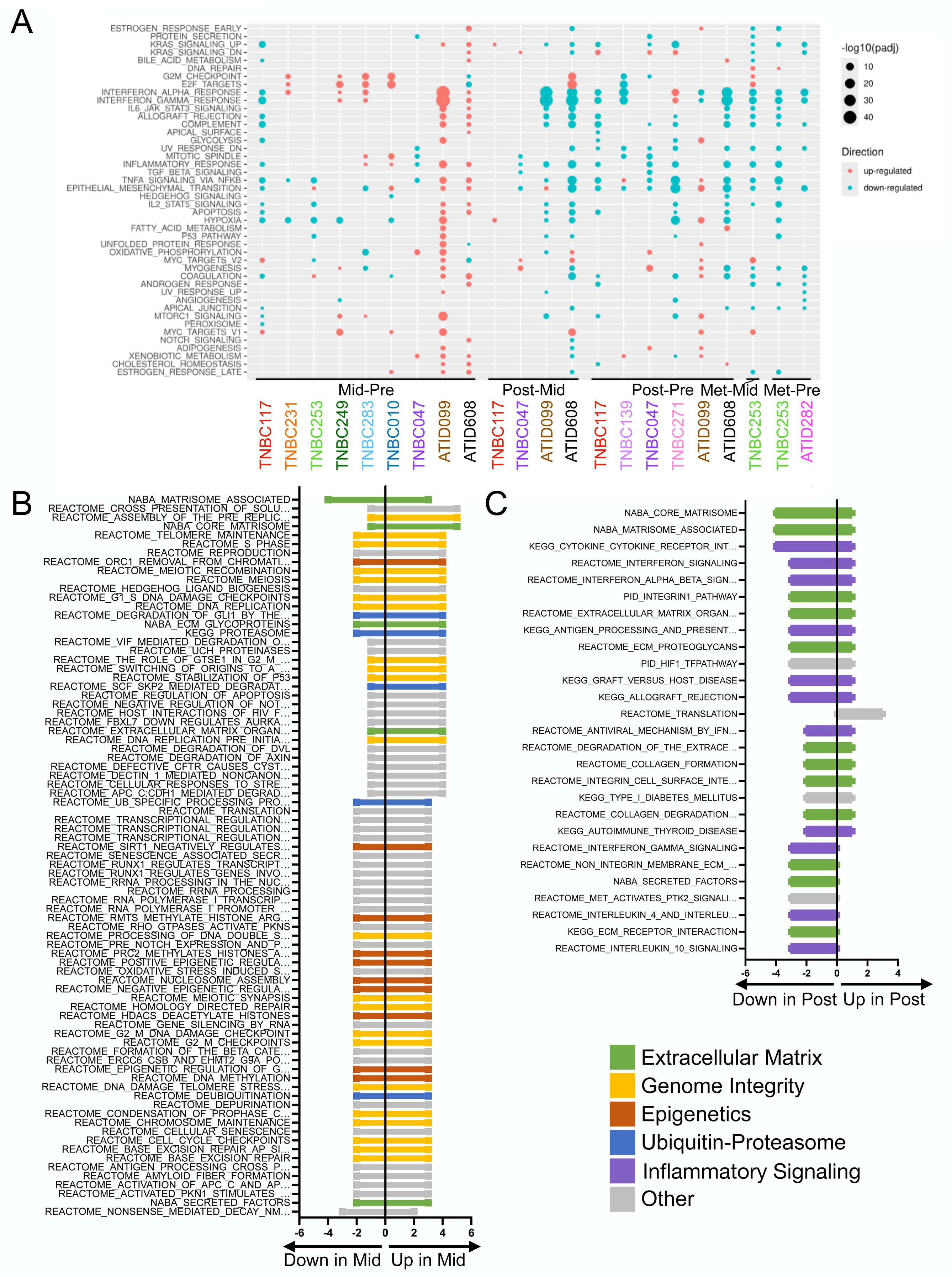
Analysis of the transcriptomes of longitudinal PDX tumor sets. (A) Pairwise comparisons of differentially regulated Hallmark pathways within the longitudinal sets. Models are indicated by patient ID with color scheme. Dot size represents significance, while color indicates the direction of change. (B) Number of models that up- or down-regulated each pathway when comparing mid-NACT to pre-NACT models. (C) Number of models that up- or down-regulated each pathway when comparing post-NACT to pre-NACT models. Color scheme indicates biological category for each pathway.

Comparing post-NACT models to their pre-NACT or mid-NACT counterparts, we observed downregulation of multiple Hallmark pathways, many of which were related to inflammatory signaling (interferon alpha and gamma responses, inflammatory response, TNFɑ signaling via NF-κB). Other commonly downregulated pathways included epithelial-mesenchymal transition and UV response down (Figure 6A). A caveat to comparisons with the post-NACT models is that they were derived after different treatment courses, including targeted therapies. Due to the limited representation of post-NACT models from each of the associated phase II trials, we cannot draw conclusions about distinct responses to each therapy. As a result, we treated these models as a single group in our analyses. Similar changes were observed in metastasis-derived models relative to their pre-NACT models. Using the MSigDB C2 pathway set for post-versus pre-NACT tumor comparisons, we identified 27 pathways significantly altered in at least 3 out of the 6 comparisons, primarily associated with ECM or inflammatory signaling (Figure 6C). We examined how these transcriptional changes impacted subtype correlation, both as categorical (Table 1) or continuous variables (Figure S7), finding instances of both subtype maintenance and subtype switching with treatment.

**Table 1:** Molecular subtyping of longitudinal patient and PDX tumors.

| **Treatment** | **Patient ID** | **Sample** | **Pre** | **Mid** | **Post** | **Met** | **Change over time** |
| --- | --- | --- | --- | --- | --- | --- | --- |
| Paclitaxel | TNBC139 | Patient | BL2 | - | UNS | - | Yes |
|  |  | PDX | UNS | - | BL2 | - | Yes |
|  | ATID282 | Patient | UNS | - | - | BL2 | Yes |
|  |  | PDX | BL2 | - | - | BL2 | No |
| Atezolizumab | TNBC117 | Patient | BL1 | BL1 | BL1 | - | No |
|  |  | PDX | BL1 | BL1 | BL1 | - | No |
|  | TNBC249 | Patient | BL2 | BL2 | - | - | No |
|  |  | PDX | BL2 | UNS | - | - | Yes |
|  | TNBC283 | Patient | BL1 | BL1 | - | - | No |
|  |  | PDX | BL1 | M | - | - | Yes |
|  | TNBC047 | Patient | N/A | N/A | N/A | - | N/A |
|  |  | PDX | BL2 | LAR | M | - | Yes |
|  | TNBC271 | Patient | N/A | - | N/A | - | N/A |
|  |  | PDX | M | - | M | - | No |
| Panitumumab<br>Carboplatin | TNBC253 | Patient | UNS | M | - | M | Yes |
|  |  | PDX | BL2 | UNS | - | M | Yes |
|  | TNBC010 | Patient | M | M | - | - | No |
|  |  | PDX | UNS | UNS | - | - | No |
|  | ATID608 | Patient | LAR | UNS | BL1 | - | Yes |
|  |  | PDX | BL1 | BL1 | BL1 | - | No |
| Everolimus<br>Bevacizumab | TNBC231 | Patient | UNS | UNS | - | - | No |
|  |  | PDX | UNS | UNS | - | - | No |
| Enzalutamide | ATID099 | Patient | BL2 | LAR | LAR | - | Yes |
|  |  | PDX | UNS | UNS | UNS | - | No |

Additionally, we used reverse-phase protein arrays (RPPA) to analyze proteomic changes in a subset of the longitudinal PDX sets. Clustering of the RPPA profiles showed that all but one of the matched sets clustered together, reinforcing the significant patient-level heterogeneity we consistently observed across these models and in TNBC generally (Figure 6A, Table S4). We also investigated protein markers associated with subtype coefficients and found the BL1 and BL2 subtypes were inversely correlated with many markers (Figure 6B). We observed associations with growth factor signaling in BL2 tumors and PI3K signaling in LAR tumors, consistent with other studies^41^.

## DISCUSSION

Here, we report the establishment of an annotated biobank of TNBC PDX models, generated from tumor tissue obtained in the context of the ARTEMIS neoadjuvant clinical trial, where patient response to therapy is known. The biobank consists of 92 PDX models derived from the tumors of 75 patients, including 12 longitudinal sets established from serially sampled tumors across trial timepoints and metastatic lesions. Most published PDX collections have been established using single biopsies obtained from the primary and metastatic tumors of patients who had undergone therapy^9,10,12,43–46^. In contrast, our collection was established from pooled fine needle aspirates (FNAs) and features 71 treatment-naïve models. Combining FNAs from different regions of the tumor is expected to capture the heterogeneity of a TNBC tumor more fully than a single biopsy can. Additionally, FNAs are less painful to patients than repeated core biopsies, increasing the likelihood that patients will consent to serial sampling. This approach enabled us to generate 12 longitudinal PDX models from serial patient sampling throughout NACT, as well as from metastatic disease.

Another distinctive feature of our biobank is the acquisition of tumor tissue alongside a single neoadjuvant clinical trial, where detailed information on patient characteristics, tumor features, and pathologic response outcomes are known. PDX models were established from patients with long-term follow-up, providing valuable clinical insight into tumor aggressiveness and sensitivity to various systemic treatments. Furthermore, nearly all patient tumors have been characterized using WES and RNA-seq, allowing for direct comparisons to their corresponding PDX tumors. Comparative analyses demonstrated that PDX tumors largely recapitulated the genomic and transcriptomic characteristics of their corresponding patient tumors and reflect the molecular and phenotypic heterogeneity of TNBC. The greatest differences in the transcriptomes between patient and PDX tumors were those related to immune and stroma interactions, which is expected when adapting human tumors for growth in immune-deficient murine hosts. Nonetheless, we saw a strong recapitulation of gene expression patterns related to signaling pathways, suggesting that these PDX models effectively capture many of the key biological features of the corresponding patient tumors. The collection includes representation from each transcriptional subtype (BL1, BL2, LAR, and M), demonstrating that the PDX biobank represents the extensive inter-patient heterogeneity characteristic of TNBC.

While we successfully established PDX models from patients who achieved pCR or RCB-I following NACT, these models did not respond with full tumor elimination in AC treatment preclinical studies *in vivo*^38^. This could be due in part to the inability to reach human-equivalent doses of chemotherapy in mice due to toxicity. Additionally, it is possible that an intact immune system, which is absent in NOD/SCID mice, is necessary to achieve this level of response^47^. Indeed, analysis of the ARTEMIS patient samples found that nearly all immune cold tumors were chemoresistant, underscoring the interplay between the immune system and chemotherapy response (Seth et al*., in preparation*). With the increasing utilization of immune checkpoint inhibitors in the neoadjuvant setting, it will be important to complement PDX studies with immunocompetent models and primary patient tumors whenever possible. Nonetheless, given that most immune cold and a portion of immune hot tumors do not respond to immunotherapy, PDX models will be particularly useful in identifying intrinsic mechanisms of therapy resistance in TNBC.

Models derived from treatment-naïve tumors provide the opportunity to investigate tumors that have not undergone therapy-induced selection, providing a valuable platform to identify potential therapies for targeting resistance in primary, treatment-naïve TNBC. A high-throughput drug screen of 16 treatment-naïve models from our collection revealed that a kinesin spindle protein inhibitor (KSPi) was broadly active across all molecular subtypes^48^. The KSPi targets the mitotic spindle through mechanisms that are independent of microtubule stability and showed efficacy in PDX models that were resistant to paclitaxel, a microtubule inhibitor used in standard TNBC treatment. In addition, subtype selectivity was observed with PRIMA-1^MET^ which exhibited enhanced efficacy in mesenchymal subtype tumors^48^.

The longitudinal models are another unique feature of our PDX collection. These models facilitate studies on how NACT impacts TNBC heterogeneity. The subclonal architecture of tumors within the longitudinal sets remained largely unchanged in response to AC, which is not unexpected given that these tumors came from patients whose tumors were largely resistant to AC (RCB-II and RCB-III). These results contrast with some post-NACT tumors and metastases where changes in subclonal composition were observed and are consistent with barcoding studies investigating chemoresistance and metastasis conducted in PDX tumors^22,38^.

Throughout our studies, we found that patient identity had a greater impact on transcriptional state than treatment timepoint or resistance status in the longitudinal PDX models. A recent study of isogenic carboplatin-sensitive and -resistant PDX models reached a similar conclusion^49^. Analysis of transcriptional changes in longitudinal tumors identified several pathways that were dysregulated in response to treatment. Inflammatory signaling was commonly downregulated in post-NACT and metastatic tumors relative to pre-and mid-NACT tumors. This finding is consistent with therapy-resistant tumors being able to evade immune recognition, perhaps by suppressing the immune system. Accordingly, immunotherapy has been a recent addition to TNBC treatment paradigms, aiming to engage the immune system in tumor elimination^2,5^.

The ubiquitin-proteasome pathway was dysregulated in mid-NACT models relative to treatment-naïve models. A previously reported genome-wide siRNA screen identified a dependence on proteasome activity in TNBC^50^. Although bortezomib, a proteasome inhibitor, did not show efficacy as a single agent in patients with metastatic breast cancer^51,52^, some responses were observed when bortezomib was combined with capecitabine^53^. We conducted a high throughput drug screen on tumors in our longitudinal collection and identified increased sensitivity to inhibitors targeting both the proteasome and neddylation pathways in mid-/post-NACT tumors compared with pre-NACT tumors^54^. Neddylation is essential for full activity of the Cullin-RING ligase complexes and thus is directly tied to the function of the ubiquitin-proteasome system^55^. Pevonedistat, a neddylation inhibitor, showed efficacy in several of the longitudinal PDX models^54^. There continues to be broad interest in investigating additional inhibitors that impact the ubiquitin-proteasome system in breast cancer, including those that target NRF1, UBA1, and PIM kinase^56–58^. Other pathways commonly dysregulated in mid-NACT models relative to pre-NACT models included genome integrity and epigenetics. Drugs targeting proteins in these pathways are also under investigation for the treatment of chemoresistant TNBC^59–66^.

In summary, our PDX collection provides a valuable resource for the TNBC community, as it recapitulates the heterogeneity characteristic of TNBC and can be used as an experimental model to address the most pressing issues facing the clinical management of TNBC, namely chemoresistance and metastasis.

## Resource Availability

### Lead Contact

Further information and requests for resources and reagents should be directed to and will be fulfilled by the lead contact, Helen Piwnica-Worms

### Materials Availability

All PDX models generated in this study are available from the authors upon request.

### Data and Code Availability

Processed RNA-seq data are available at GEO #######. Additionally, clinical and genomics data characterizing the PDX biobank is available at pdxportal.research.bcm.edu. Code used to parse, analyze, and visualize the genomics data is available at ####.

## Acknowledgements

We are grateful to the patients who provided tumors for PDX model establishment. We thank Dr. Sendurai Mani, Dr. Abena Redwood, Dr. Yan Jiang, Aaron McCoy, and Yizheng Tu for contributions in the early stages of PDX establishment. We thank Drs. Naoto Ueno, Jennifer Litton, Bora Lim, and Senthil Damodaran for their leadership as PIs of the post-AC Phase II clinical trial arms for ARTEMIS. Drs. Rosalind Candelaria, Lumarie Santiago, Beatriz E. Adrada, Deanna L. Lane, and Wei T. Yang facilitated tumor acquisition for PDX establishment. PDX models were established through a generous gift from the Cazalot family and from funds from the MD Anderson Cancer Center Breast Cancer Moon Shot Program. Additional funding sources that supported this work include the Cancer Prevention and Research Institute of Texas RP150148 (to H.P.-W.), RP160710 (to W.F.S., H.P-W., and J.T.C.). This study was funded in part by generous philanthropic contributions to The University of Texas MD Anderson Cancer Center Moon Shots Program, a Conquer Cancer Career Development Award supported by Fleur Fairman (to C.Y.), philanthropic support from the Still Water Foundation (to C.Y. and S.L.M.), and the Winterhof Foundation (S.L.M.). Dr. Yam was additionally supported by the 2018 Gianni Bonadonna Breast Cancer Research Fellowship (Conquer Cancer Foundation), the Allison and Brian Grove Endowed Fellowship for Breast Medical Oncology, and the Susan Papizan Dolan Fellowship in Breast Oncology. The MD Anderson Research Histology Core Laboratory, the Advanced Technology Genomics Core, and the Characterized Cell Line Core are supported in part by NCI grant P30CA016672. The Functional Proteomics Reverse Phase Protein Array Core was supported in part by The University of Texas MD Anderson Cancer Center, P30CA016672, and R50CA221675.

## Author contributions

**Conceptualization:** ALR, JTC, HPW

**Methodology:** ALR, EP, PdH, GVE, WFS, AMT, SLM, JTC, HPW

**Formal Analysis:** ALR, YQ, JTC

**Investigation:** SC, JSS, SJJ, XZ, CF, RL

**Resources:** EER, JBW, LH, GMR, BA, CY, SLM

**Data Curation:** FH, EER, JBW, RG, CY, YQ, JTC

**Writing – Original Draft:** ALR, YQ, JTC, HPW

**Writing – Review and Editing:** ALR, GVE, AMT, WFS, LH, SLM, JTC, HPW

**Visualization:** ALR, YQ

**Supervision:** WFS, SLM, JTC, HPW

**Project Administration:** ALR, FH, EER, JBW, RG, GMR, BA, CY, WFS, SLM, JTC, HPW

**Funding Acquisition:** WFS, AMT, SLM, JTC, HPW

## Declaration of interests

S.L.M. and E.E.R. are employees of and hold stock in Eli Lilly and Company. The spouse of A.M.T. works for Eli Lilly and Company. W.F.S. owns founder stock in Delphi Diagnostics and publicly traded stock in IONIS Pharmaceuticals and Eiger BioPharmaceuticals; is a consultant/advisor to Merck, Astra Zeneca. W.F.S. and C.F. are co-inventors of pending patent “Targeted Measure of Transcriptional Activity Related to Hormone Receptors”, United States, Provisional Patent Application Serial No. 62/329,774. G.V.E. receives sponsored research funding from Chimerix, Inc. and experimental compounds from the Lead Discovery Center of Germany. C.Y. has received research funding (to the institution) from Genentech, Gilead, BostonGene, Sanofi, Amgen, Pfizer, Astellas, and Novartis and has served on advisory boards for Gilead and Cytodyn. The remaining authors declare no competing interests.

## Supplemental Information

Document S1. Figures S1-S8 and Tables S2 and S4

Table S1. Excel file containing characteristic information for each PDX model which is too large to fit in PDF, related to Figure 1.

Table S4. Excel file containing RPPA data which is too large to fit in PDF, related to Figure S8.

## METHODS

### Collection of patient-derived materials

All research conducted in human patients followed national guidelines including the Health Insurance Portability and Accountability Act (HIPAA) privacy and security rules^67^ and the Common Rule (http://www.hhs.gov/ohrp/humansubjects/commonrule/). Informed consent was obtained from all patients who provided samples for PDX model establishment. Patients were enrolled in the ARTEMIS trial (NCT02276443), an MD Anderson Cancer Center IRB-approved protocol (2014-0185).

### Establishment of PDX collection

All experimental procedures were approved by the Institutional Animal Care and Use Committee (IACUC) at MD Anderson Cancer Center under IACUC protocol 00000978-RN01. Ends for animal experiments were selected in accordance with IACUC approved criteria. Female NOD/SCID mice (NOD.CB17-*Prkdc^scid^*/NcrCrl) were obtained from Charles River, National Cancer Institute colony.

PDX models were established according to published protocols^8,14^. Three to four weeks prior to tumor cell engraftment, the fourth mammary fat pads (MFPs) of 4- to 5-week-old NOD/SCID female mice were pre-humanized with green fluorescent protein (GFP)–labeled immortalized human mammary stromal fibroblasts (EG cells). FNA biopsies obtained in the clinic were immediately placed in DMEM/F12 media (HyClone Cat# SH30023.01) with 1x antibiotic/antimycotic (Corning cat # MT30004CI) and maintained on ice during transport to the laboratory. Enzymatic digestion of the samples used complete DMEM/F12 media supplemented with 3mg/mL collagenase (Roche cat# 1088793) and 250 U/mL of hyaluronidase (Sigma cat# H-3506) with incubation at 37°C in a rotator for 30 to 90 minutes. Following digestion, samples were washed with media and incubated with red blood cell lysis buffer (Sigma Cat# R7757). Cell pellets (passage 0, P0 cells) were washed with media and resuspended in DMEM/F12 supplemented in 5% bovine calf serum (BCS) and Matrigel (50/50; Corning) before injection into the right fourth MFPs of mice. Tumor cells were injected in a total volume of 30-50 µL. On average, 4,000-100,000 cells were implanted into two mice for the first PDX generation, depending on the tumor cell yield. Typically, cells were implanted into mice within two hours of the FNA biopsy reaching the laboratory.

Engrafted mice were monitored for up to 6 months for tumor growth, at which point mice without palpable tumors were euthanized. When PDX tumors reached approximately 1000 mm^3^, they were collected and stored as snap-frozen tumor pieces, dissociated single cell suspensions, or formalin-fixed, paraffin-embedded tissue blocks. The dissociation of PDX tumors was the same as for FNA biopsy processing, with 3 to 5 h digestion times. P1 tumor cells were immediately implanted into the next generation of mice (n=2), which were not pre-humanized with stromal fibroblasts. Tumors were passaged through a total of three generations of mice, with 6 mice used for each subsequent generation. PDX models that reached P3 (the 3rd generation of mice) and were validated as described below were considered established.

### Quality control of PDX models

Genomic DNA (gDNA) was extracted from PDX tumor cells using the DNeasy Blood & Tissue Kit (Qiagen). We used three approaches to validate the identity of each PDX model throughout passaging in mice^14,22^. First, we determined the human origin of each PDX tumor by quantifying the ratio of human-to-mouse gDNA in PDX tumors via qPCR with a human RNaseP gene probe (20x human RNaseP copy number assay, FAM-TAMRA, Life Technologies) and a mouse Trfc gene probe (20x mouse Trfc copy number assay, VIC-TAMRA, Life Technologies). The absolute calibrators were gDNA from TaqMan™ Control Human Genomic DNA (ThermoFisher) and from Mouse Genomic DNA (Promega). qPCR was conducted using the 2x TaqMan gene expression master mix (Life Technologies). The relative ratio of human and mouse gDNA in each tumor sample was calculated using the ΔΔCt method as described previously^68^. PDX tumors consistently exhibiting <10% human genomic DNA were discarded and not included in any subsequent analyses.

Second, to confirm clearance of the GFP-labeled EG cells used to pre-humanize the first generation of mice, PCR was performed against GFP and human GAPDH. Amplicons were visualized on agarose gel after electrophoresis. Primer pairs for GAPDH were: Fwd 5′-AAGTTCATCTGCACCACCG; Rev 5′-TCCTTGAAGAAGATGGTGCG. Primer pairs for GFP were: Fwd 5′-ACATCATCCCTGCCTCTAC; Rev 5′-TCAAAGGTGGAGGAGTGG. All PDX models herein were confirmed to be GFP negative by this test.

Lastly, to establish a DNA ‘fingerprint’ for each PDX model for future validation purposes, short-tandem repeat (STR) DNA fingerprinting was conducted using 50 ng of gDNA through the MDACC Characterized Cell Line Core. The STR profiles from Promega 16 High Sensitivity STR Kit (Catalog# DC2100) analyses were compared to commercial databases (DSMZ/ATCC/JCRB/RIKEN) of approximately 2500 known profiles, as well as the MDACC Characterized Cell Line Core database of approximately 2000 known DNA fingerprint profiles. Each PDX tumor had a unique STR profile, confirming that none were contaminated with human cell lines and that cross-contamination among the models did not occur. All tumors within each longitudinal set shared the same STR profile, indicating they were derived from the same patient. The matching profiles of subsequent PDX passages demonstrated consistent identities.

### Histology

Patient and PDX tumor biopsies stored as formalin-fixed paraffin embedded tissue blocks were cut into 5 μm sections and analyzed by hematoxylin and eosin (H & E) staining. Histologic properties were evaluated by a breast pathologist (Dr. L. Huo). Immunohistochemical staining was performed using standard methods. After deparaffinization of the sections with xylene and rehydration via graded aqueous-ethanol baths, antigen retrieval was performed using Reveal decloaker solution (Biocare Medical, RV1000M) and an EX-Retriever v.3.0 microwave. Endogenous peroxidases were quenched by treatment with Dako Dual Enzyme block (S2003). The human-specific mitochondria antibody (Abcam ab92824) was diluted 1:800 and the pan-cytokeratin antibody was diluted 1:100 (Abcam ab86734). The M.O.M kit (Vector Laboratories MP-2400) was used to develop staining for both antibodies.

### CBR-luciferase labeling of PIM056

To generate a sub-line of PIM056 expressing fluorescent and bioluminescent markers, we followed previously established protocols^22,69^. Briefly, a PIM056 tumor was harvested, dissociated into single cells, and depleted of mouse cells using magnetic-activated cell sorting (Miltenyi, Cat. # 130-104-694). Two million human tumor cells were cultured under mammosphere conditions using 5 mL of complete Mammocult medium (StemCell Technologies) in ultra-low attachment plates (Corning) in the presence of recombinant lentivirus (approximate MOI = 8) encoding Click beetle red luciferase (CBRLuc) and mCherry, followed by adding 10µg/mL polybrene (Sigma) twelve hours later. Twenty-four hours after transduction, the media was refreshed. Four days after lentiviral transduction, cells were pelleted, organoids were dissociated with TrypLE Express dissociation reagent (Gibco), washed, and resuspended in PBS containing 0.5% BSA. Cells were stained with Sytox Blue viability dye (Life Technologies), then sorted on a BD Aria Fusion fluorescence activated cell sorter (FACS) to collect viable, mCherry-positive cells. These cells were washed with media and engrafted into the MFP of one NOD/SCID mouse as described above. Once the tumor reached 1000 mm^3^, it was harvested, dissociated into single cells and immediately transplanted into a second generation of NOD/SCID mice. This process was repeated on second passage CBRLuc-labeled tumor cells. Aliquots of cells were stored in cryopreservation media and frozen after each passage.

### Bioluminescence imaging (BLI)

BLI was performed as previously described^21^. Animals received an intraperitoneal injection of D-luciferin (150 µg/g body weight, GoldBio) in phosphate-buffered saline. Ten minutes after injection, animals were anesthetized via isoflurane and imaged with a charge-coupled device camera-based BLI system (IVIS Spectrum, PerkinElmer). Signals were displayed as photons/s/cm^2^/sr for image representation. To detect distal organ metastases, tissues were assessed with BLI *ex vivo* at necropsy. Lungs, liver, brain, and bone were subjected to BLI individually.

### Sequencing patient samples and early-passage PDX tumors

Initially, samples were sequenced by WES at the Cancer Genetics Laboratory at MD Anderson Cancer Center. Genomic libraries from approximately 500 ng gDNA were prepared using the standard KAPA paired-end sample preparation kit (KAPA Biosystems) according to the manufacturer’s instructions. SureSelect Human All Exon Kit, version 4 (Agilent Technologies) was used to enrich sequencing libraries for exomes. Samples were pooled 2 per lane and paired end 2 × 75 bp sequencing was performed using the Illumina HiSeq 2000. Later in the study, sequencing was performed by the Sequencing and Microarray Facility at MD Anderson Cancer Center. They followed a similar protocol as described above and also used the Agilent SureSelect Human All Exon Kit, version 4 to enrich for exomes. In contrast, they used the Agilent SureSelectXT Reagent Kit (Agilent Technologies) for library preparation and sequenced on an Illumina HiSeq4000 sequencer.

### Whole-exome sequencing data analysis

We processed the NGS data using the BETSY system^70^. Reads from stromal mouse sequences were identified and discarded using Xenome^71^. We aligned the remaining non-mouse reads to the human reference (hg19) following the GATK best practices^72,73^. Bam files were sorted and indexed using samtools^74^. Duplicates were identified using Picard tools^73^ and indels were realigned using GATK.

### Somatic mutation calls

For somatic variant callers, we used the patient’s blood sample as the germline reference when available. For unavailable germline samples, we generated a common normal using 213 blood germline samples from the ARTEMIS trial that were each downsampled to 1% of reads and merged. We set minPruning = 3 in MuTect2 to accelerate computation. We also applied a custom filter designed to identify alignment artifacts from repeat regions^22,75^. To identify somatic variants, we employed a consensus approach^76,77^ consisting of three algorithms to call SNPs and short and long indels (MuSE^78^, MuTect2, and Varscan2^79^). Variants identified by two out of the three algorithms were considered true variant calls. We annotated the variants using Annovar^80^ and SnpEff^81^. We further applied a quality filter of a minimal alternative allele of 5. All mutations that passed this filter were used in downstream analysis.

### Copy number assignments

Analysis of copy number alteration for paired tumor-normal whole exome sequencing samples was performed using FACETS^82^. To detect major changes, the critical value for segmentation was set to 100, and minimum number of heterozygous SNPs in a segment used for bivariate t-statistic during clustering was set to 30. To reduce hyper-segmentation, scanning window size was set to 250 bp. Other parameters remain the same as default values. From the segmentation files produced by FACETS, we extracted the estimated total copy number (under column tcn.em) and plotted the values across the human genome as a heatmap. Plots of longitudinal PDX sets derived from the same patient were grouped together.

### PyClone

To select variants for subclone analysis, we discarded those with low (<20) or high (>1000) read depth coverage, low alternative allele reads (<5), or frequency (<5%) in any samples, and those that appeared to be errors by manual inspection of the alignments. We discarded variants with high total copy number (>6), and those that occurred in regions where the copy number varied across samples. Using the read count, copy number, and purity prediction for the selected variants, we used PyClone^39^ to estimate the cancer cell frequencies (CCFs). We used the runanalysispipeline script in Pyclone (v0.13.1) under the default settings: basemeasureparams (alpha = 1; beta = 1), betabinomialprecisionparams (prior rate = 0.001; shape = 1; proposal precision = 0.01; value = 1000), concentration (prior rate = 0.001; shape = 1; value = 1), density = pyclone_ betabinomial, numiters = 10,000, and errorrates = 0.001, except for prior for method to set the possible genotypes = totalcopynumber, for all samples. Output statistics and plots were generated after 1,000 iterations as burn-in.

PyClone analysis identified multiple clusters comprising one or more variants. We noticed that some clusters were unstable and were comprised of variants whose cluster assignments were unstable throughout the MCMC sampling. To quantify the unstable clusters, we calculated a pairwise matrix containing the probabilities that each pair of variants were assigned to the same cluster across MCMC samples. We then calculated a matrix containing the difference of probabilities of a variant with other variants within its cluster and outside its cluster. We accepted a cluster as robust if the number of variants in the cluster is greater than 5 and the median difference of probabilities is greater than 80%.

### Mutation consistency rate between PDX and patient

Consistency rate between corresponding PDX and patient samples was defined as the number of common mutations in both a PDX tumor and corresponding patient sample divided by the number of all mutations in the patient sample. The range of consistency rate is 0 to 1.

### Frequently mutated genes

Mutation frequency of a variant was calculated as patient-based, not sample-based. If a variant occurred in any sample belonging to the same patient, it was considered to occur once in that patient. Silent variants, as well as those falling into intergenic and intronic regions, were removed from this analysis. The ClinVar database^83^ was downloaded and processed to obtain a collection of pathogenic mutations. Variants in our dataset that mapped to this collection were considered pathogenic.

### Tumor mutation burden analysis

Mutation burden for each sample was calculated as the number of nonsynonymous mutations divided by the sequenced length. Here, nonsynonymous mutations include all those mutations that lead to a change in protein sequence, e.g., missense, startloss, stopgain, stoploss.

### Mutation signature analysis

Mutation signature analysis was carried out using MuSiCal v1.0^84^ by refitting the trinucleotide profile in our dataset to the program’s Breast.AdenoCA signature collection. The resulting exposure scores of each signature were scaled to sum up to 100% for each sample. Signatures with less than 15% exposure in all PDX models were considered as “others” when plotting.

### RNA sequencing data analysis

We processed the data using pipelines implemented in the BETSY system^70^. We aligned the fastq reads to human genome hg19 using STAR^85^. Gene-level read counts were calculated with HTSeq-Count^86^ and transcripts per million (TPM) were summarized using RSEM^87^. Batch effect correction of the original TPM was carried out using ComBat^88^. The batch-corrected TPM values were used in downstream analyses, unless stated otherwise.

### TNBC subtyping

Vanderbilt TNBC subtypes^34,89^ were assigned using the TNBCtype public server: https://cbc.app.vumc.org/tnbc/. The assignments were carried out by sequencing batch. Original TPM values from samples within the same sequencing batch were submitted to the server for simultaneous assignment, minimizing potential issues related to normalization across different batches. Two of the subtypes assigned by the server, IM and MSL, are heavily impacted by stromal infiltration^33^ and were excluded in our PDX tumor analysis. To convert these two subtypes, as well as the unstable (UNS) group, to other valid subtypes (BL1, BL2, M, LAR), we applied the following algorithm, only utilizing p-values and correlation coefficients of the four valid subtypes (BL1, BL2, M, LAR). We assigned the subtype with the largest correlation coefficient among subtypes with p-value ≤ 0.05 and correlation coefficient ≥ 0.1. The difference between this and the next largest coefficient had to be ≥ 0.05. If no subtype met this criteria, “UNS” was assigned^89^. The significance test of subtype co-occurrence was carried out using the statistical model described by Zhou et al^90^. A one-sided p-value was calculated using an implementation of the method in R.

### Gene expression comparison between corresponding patient and PDX samples

Comparisons of the gene expression profiles (nsample=84) were carried out using Spearman correlation of the 1000 most variable genes from PDX models and the corresponding genes from patient tumor samples.

### Unsupervised consensus clustering

We performed unsupervised consensus clustering using Pearson correlation distance and a hierarchical clustering algorithm by using the R package ConsensusClusterPlus^91^. The heatmap was plotted using R package ComplexHeatmap^92^.

### Single-sample pathway scores

We calculated single-sample pathway scores using the ssGSEA algorithm implemented in R package GSVA^93^.

### Pairwise pathway analyses between timepoints

We performed pairwise pathway analyses on the longitudinal sets of PDX models using the pre-ranked GSEA algorithm implemented in R package fgsea^94^ with gene expression fold change for ranking.

### Calculating conservation scores

Each patient sample-PDX model pair was scored for degree of conservation (0, 0.5, or 1) in five categories: mutations, mutation burden, subclonal architecture, gene expression, and transcriptional subtype. The total conservation score was the sum across all categories for each pair of samples. For mutations, the median conservation proportion of mutations across the dataset was 0.554. All pairs with a higher conservation proportion scored 1, any pairs between 0.300 and 0.554 scored 0.5, and any pairs below 0.300 scored 0. For mutation burden, we previously classified samples into high (>15 mutations/Mb), medium (5-15 mutations/Mb), and low categories (<5 mutations/Mb). Any pairs that matched scored 1 and pairs that did not match scored 0. For subclonal architecture, pairs with the same number of subclones, with proportions within 25% of each other, scored 1, pairs with the subclones varying >25% in proportion scored 0.5, and pairs with different numbers of subclones scored 0. For gene expression, the median Spearman correlation coefficient for matched patient-PDX samples was 0.588, while the median coefficient between any patient-PDX pair was 0.351. Paired samples with a correlation coefficient greater than the median 0.588 scored 1, pairs between 0.351 and 0.588 scored 0.5, and pairs below 0.351 scored 0. For transcriptional subtype, we previously classified samples using the 4-class TNBCtype, allowing for both a primary subtype assignment and multiple significant subtype correlations. Pairs with the same primary subtype assignment scored 1, pairs with multiple significant subtypes that shared at least one in common scored 0.5, and pairs with no subtypes in common scored 0.

### Preclinical studies in mice

NOD/SCID mice were implanted with 0.5-1.0 x 10^6^ cells in the fourth MFP. When tumors reached ∼100-150 mm^3^, mice were randomized into vehicle or chemotherapy groups for dosing. Mice received a single dose of both doxorubicin (0.5 mg/kg) and cyclophosphamide (50 mg/kg) or sterile water (vehicle) via intraperitoneal injection. Tumor volumes were monitored via caliper measurements twice weekly until allowable tumor burden was reached.

### Reverse-phase protein array (RPPA)

Tumor fragments were collected from each PDX model and flash-frozen in liquid nitrogen. Frozen tumor pieces were homogenized using a Next Advance Bullet Blender, followed by sonication in lysis buffer containing 1% Triton X-100, 50mM HEPES pH 7.4, 150mM NaCl, 1.5mM MgCl_2_,1mM EGTA, 100mM NaF, 10mM Na_4_P_2_O_7_, 1mM Na_3_VO_4_, 10% glycerol, and protease and phosphatase inhibitors. Samples were centrifuged at 13,000rpm for 10 minutes and supernatants were collected. We quantified the protein concentration by Bradford assay, then adjusted each sample to a final concentration of 1.0-1.5 µg/mL with lysis buffer. Protein lysate was mixed with 4X SDS+β-mercaptoethanol buffer and boiled for five minutes, then stored at −80°C. RPPA was then performed as previously described^95^. We used tumor pieces from three individual mice per model whenever possible, otherwise multiple tumor pieces from the same mouse were used as replicates when necessary.

Relative protein levels for each sample were determined and normalized for protein loading by algorithm SuperCurve^96^. Log2-transformed normalized values were used in downstream analysis. Samples from longitudinal PDX sets derived from the same patient were measured in the same batch. Outlier samples determined by SuperCurve loading correction factors and principal component analysis were excluded. Unsupervised clustering of RPPA expression was carried out using the top 100 most variable (by MAD) antibody markers out of 331 commonly available markers for all samples, and a heatmap of the median-centered expression values was plotted. Expression values of proteins (CDH2 and VIM) that were missing values (because they were not measured in every batch) were not included in the clustering and added to the heatmap after clustering. Correlation analysis between protein expression and Vanderbilt TNBC subtype coefficients were also performed on these 331 common markers. The top 5 most-significant markers that were positively and negatively correlated for each of the 4 subtypes (BL1, BL2, LAR, M) were selected and shown in a dot plot.

