## SupplementalMaterial for "An annotated biobank of triple negative breast cancer patient-derived xenografts featuring treatment-naïve and longitudinal samples throughout neoadjuvant chemotherapy"

**Figure S1: Histological comparison of patient and PDX samples from longitudinal sets.** H&E staining of patient (top) and PDX (bottom) tumor sections for each longitudinal set of samples. Patient identifier is shown on the left for each patient and PDX pairing. Biopsy timepoint is indicated above each column of images. Scale bars are 50μm.

**Figure S2: Association of tumor characteristics with take rate and PDX model growth (**A) Number of samples and (B) rate of successful PDX model establishment based on clinical timepoints (C) Number of samples and (D) rate of successful PDX model establishment based on patient RCB status (E) Number of samples and (F) rate of successful PDX model establishment based on two-year metastasis status of patients (G) Time to reach P3 for each successful PDX model, with longitudinal models labeled and shown in color (H) Time to reach P3 compared against transcriptional subtype of PDX model (I) Time to reach P3 compared against RCB status

**Figure S3: Copy number summary of PDX collection** Copy number calls were derived from WES data using FACETS. Amplifications and deletions are indicated by the color scale. Longitudinal sets are indicated by bars to the right of PDX IDs.

**Figure S4: Subtype coefficients for each PDX models underscores TNBC heterogeneity** Coefficients were calculated by TNBCtype for each PDX model and plotted in order of primary subtype assignment: (A) BL1, (B) BL2, (C) LAR, (D) M, and (E) UNS. Significantly correlated subtypes are noted by text below each PDX name. Longitudinal sets of models are grouped together and plotted separately (Figure S7).

**Figure S5: Summary of conservation scores between patient and PDX samples** Each patient sample-PDX model pair was scored on a scale of 0, 0.5, or 1 for conservation of features in five categories: mutations, mutation burden, subclonal architecture, gene expression, and transcriptional subtype. The overall conservation score for a sample was the sum of the scores for each category. Samples were then classified based on biopsy timepoint (A), mutation burden (B), RCB (C), patient metastatic status (D), patient subtype (E), or PDX subtype (F) and conservation scores were plotted per group. No significant differences in conservation score were observed across these categories. (G) Conservation scores plotted across the timepoints of the longitudinal PDX sets.

**Figure S6: Subclonal architecture of longitudinal patient and PDX samples** Fish plots indicate the number and prevalence of each clone or subclone at the labeled timepoint for patient and PDX samples. Corresponding patient and PDX samples are stacked vertically with patient ID indicated above, and shared colors within each pairing indicate the same subclone.

**Figure S7: Subtype coefficients for longitudinal PDX models** Coefficients were calculated by TNBCtype for each PDX model and plotted in stacked bar graphs. Each longitudal set of PDX models is graphed together. Significantly correlated subtypes are noted by text below each PDX name.

**Figure S8: RPPA analysis of matched PDX sets** (A) Heatmap of median centered z-scores for protein biomarkers clustered across longitudinal PDX models. Annotations for each sample include patient ID, timepoint, and transcriptional subtype. (B) Dot plot of the five protein markers with the strongest positive and negative correlation to each of the four transcriptional subtypes

### Figure S1

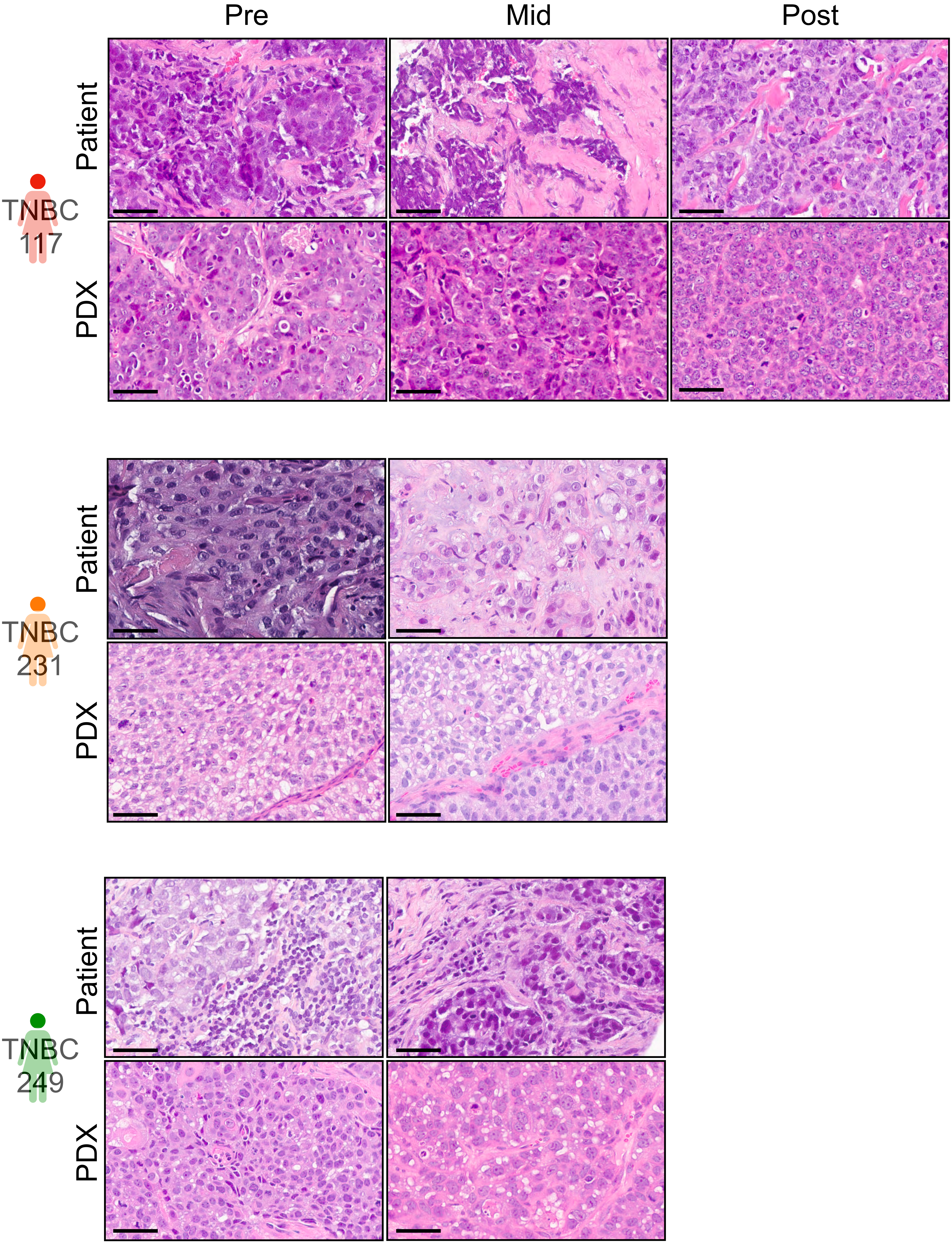
Pre
Mid
Post
Patient
PDX
TNBC 117
Patient
PDX
TNBC 231
Patient
PDX
TNBC 249

#### Figure S1 (continued)

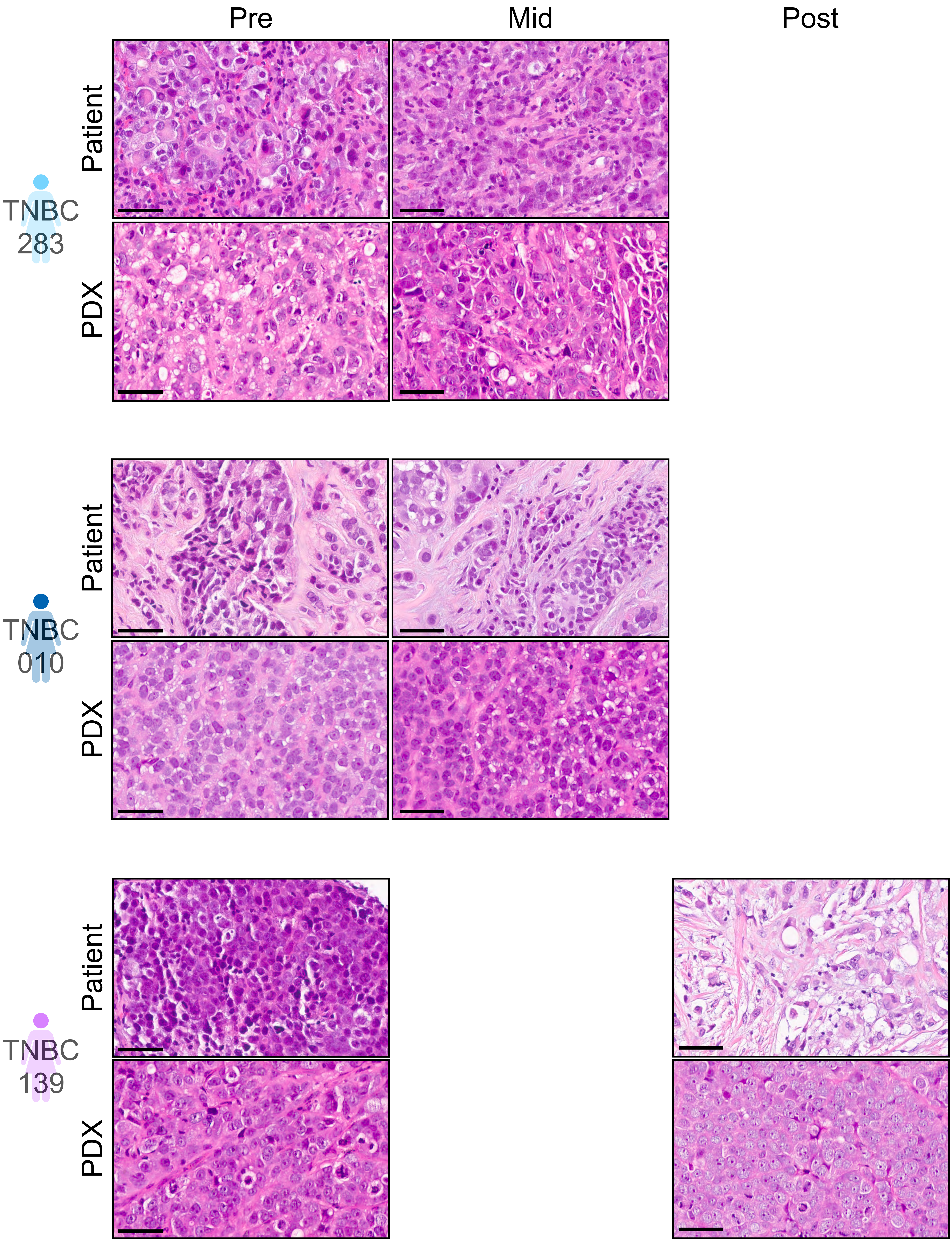

### Figure S1 (continued)

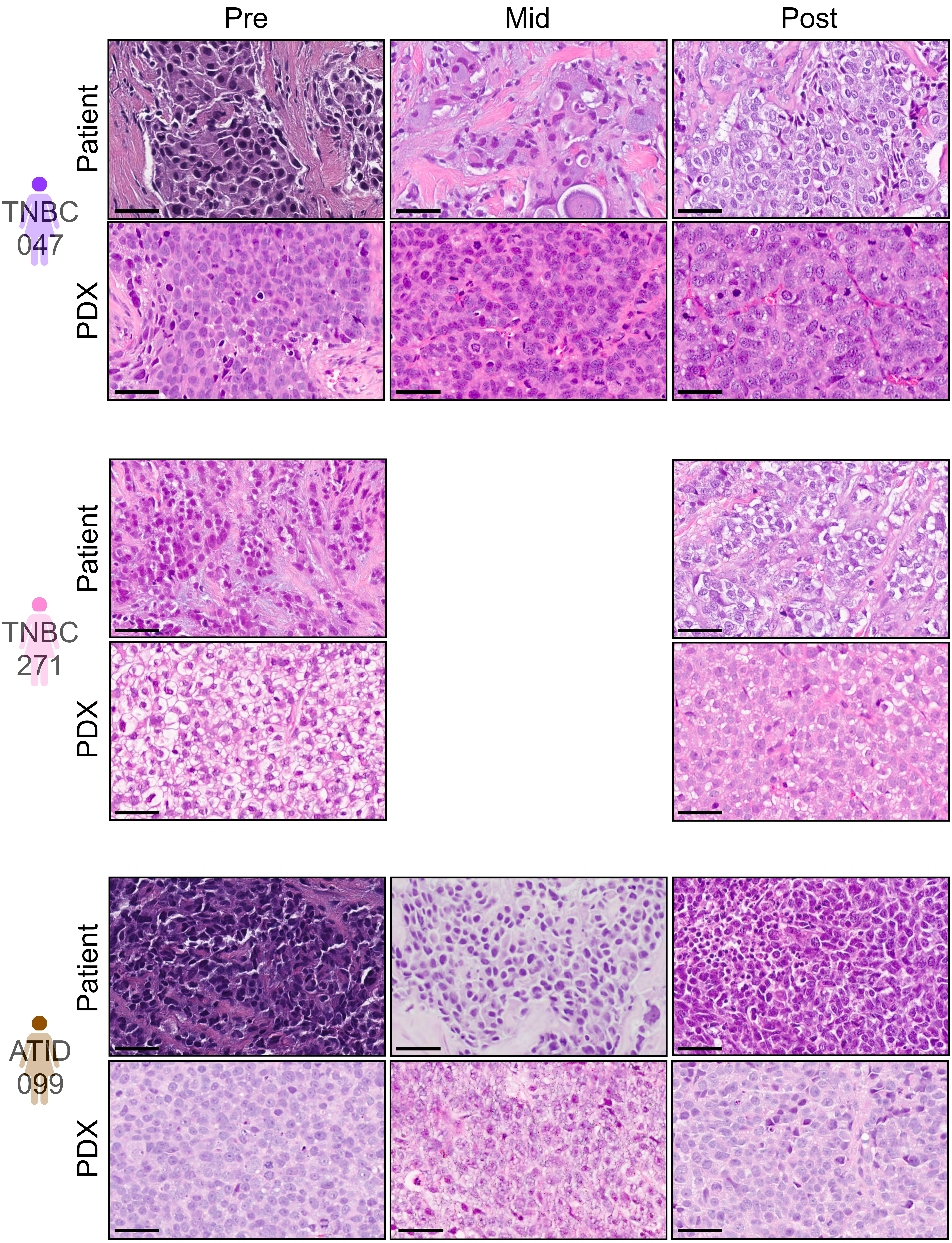

#### Figure S1 (continued)

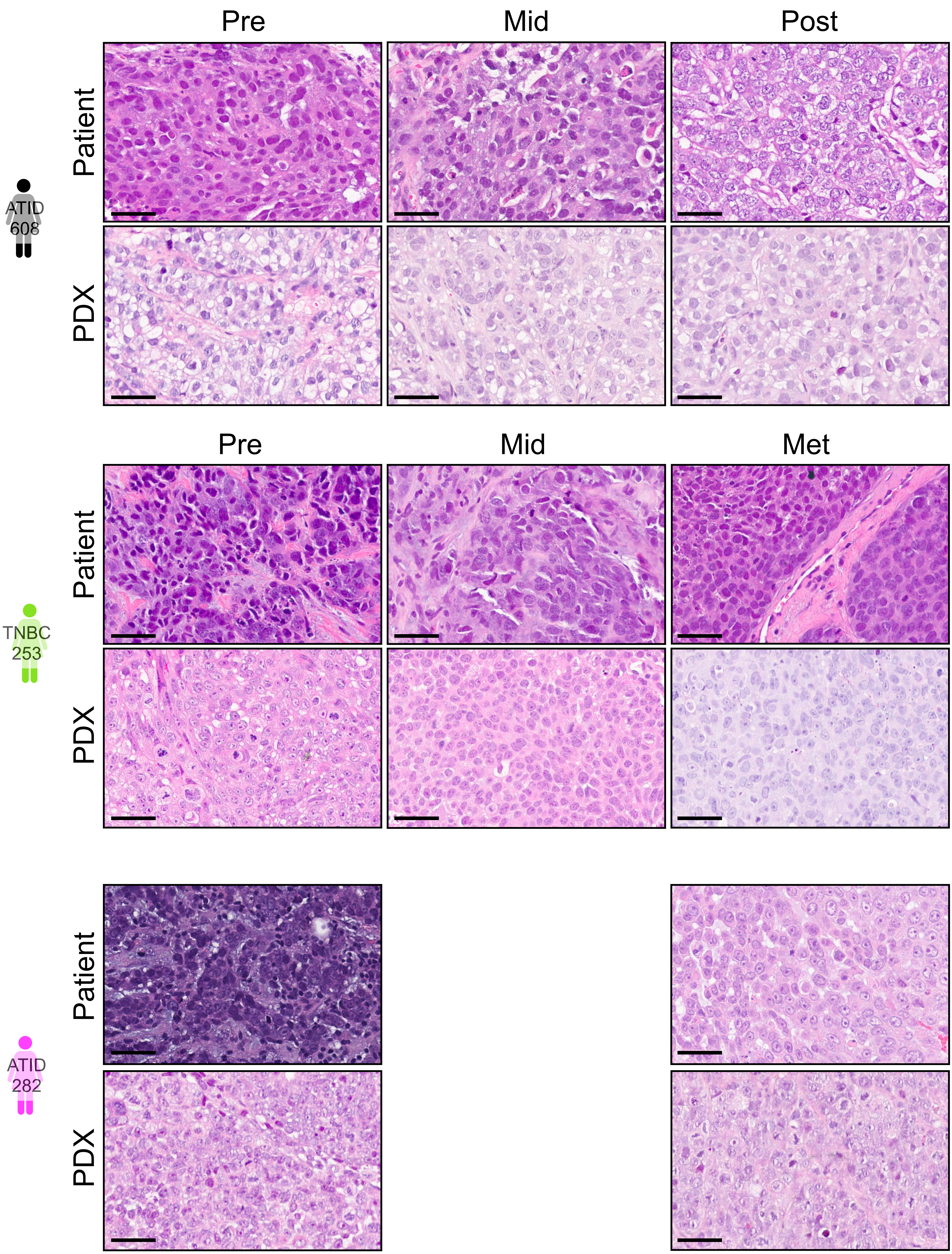

### Figure S2

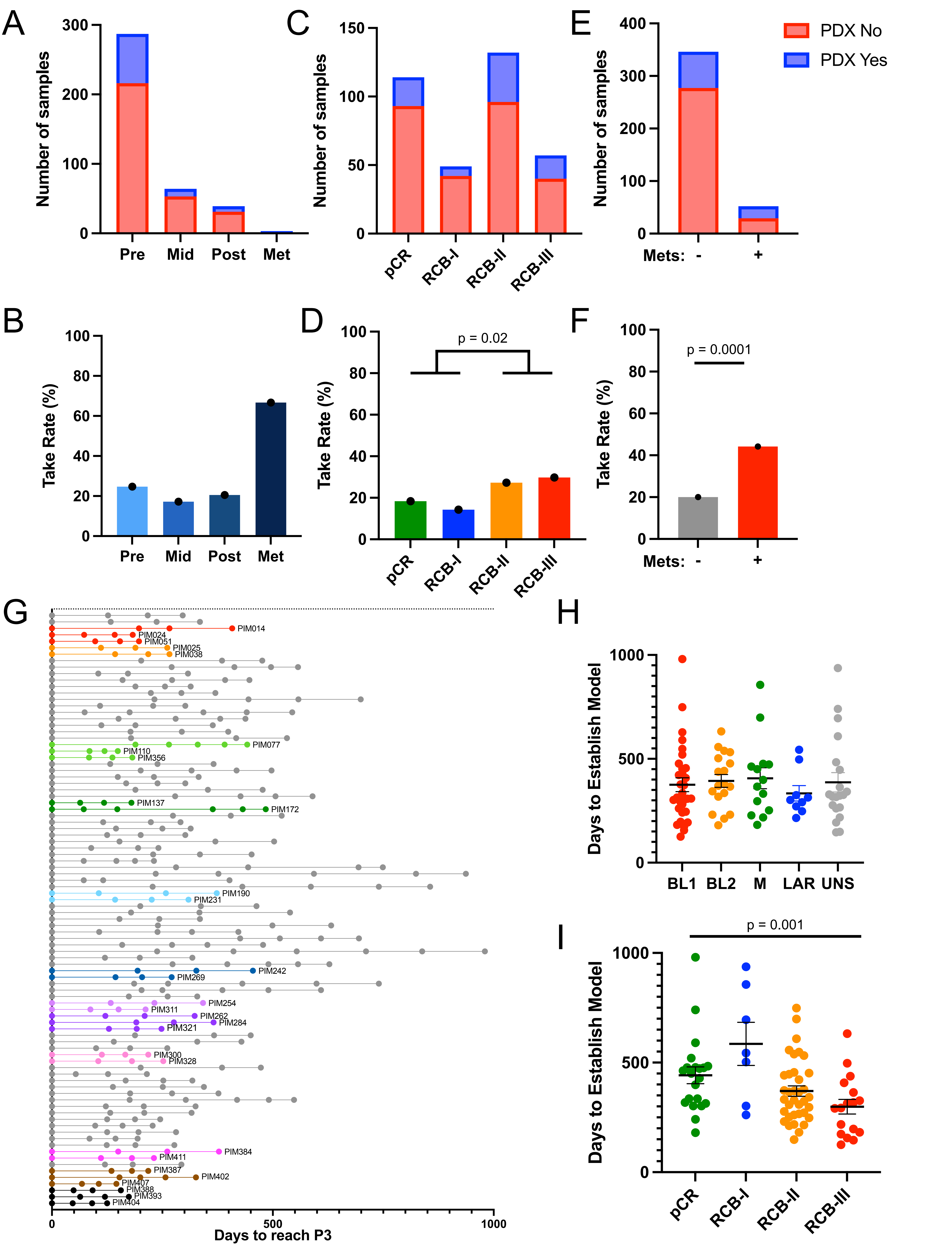
A
Number of samples
300
200
100
0
Pre
Mid
Post
Met
C
Number of samples
150
100
50
0
pCR
RCB-I
RCB-II
RCB-III
E
Number of samples
400
300
200
100
0
Mets: -
+
PDX No
PDX Yes
B
Take Rate (%)
100
80
60
40
20
0
Pre
Mid
Post
Met
D
p = 0.02
Take Rate (%)
pCR
RCB-I
RCB-II
RCB-III
F
p = 0.0001
Take Rate (%)
Mets: -
+
G
PIM014
PIM024
PIM051
PIM025
PIM038
PIM077
PIM110
PIM356
PIM137
PIM172
PIM190
PIM231
PIM242
PIM269
PIM254
PIM311
PIM262
PIM284
PIM321
PIM300
PIM328
PIM384
PIM411
PIM387
PIM402
PIM407
PIM388
PIM393
PIM404
0
500
1000
Days to reach P3
H
Days to Establish Model
1000
500
0
BL1
BL2
M
LAR
UNS
I
p = 0.001
Days to Establish Model
1000
500
0
pCR
RCB-I
RCB-II
RCB-III

### Figure S3

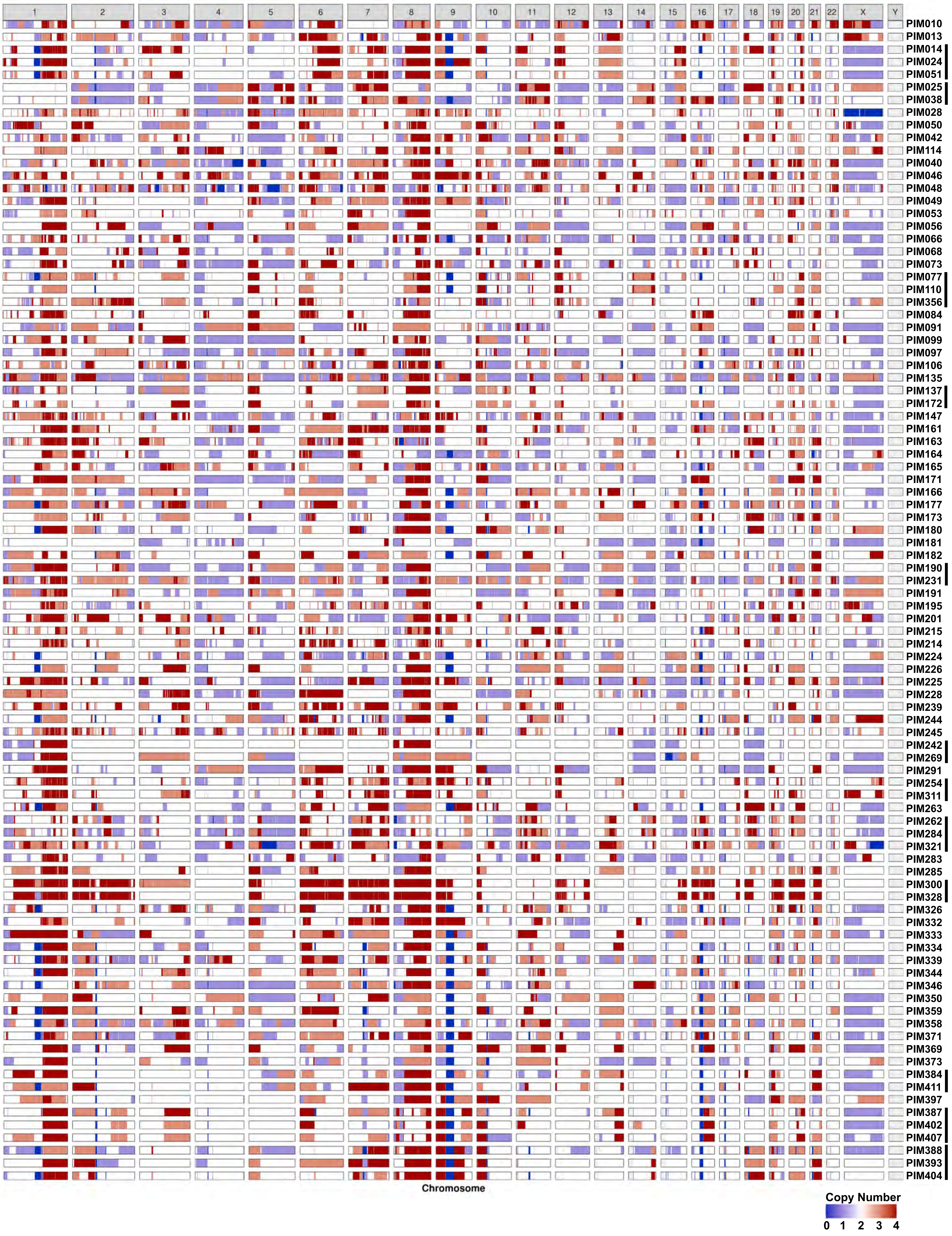
X
Y
PIM010
PIM013
PIM014
PIM024
PIM051
PIM025
PIM038
PIM028
PIM050
PIM042
PIM114
PIM040
PIM046
PIM048
PIM049
PIM053
PIM056
PIM066
PIM068
PIM073
PIM077
PIM110
PIM356
PIM084
PIM091
PIM099
PIM097
PIM106
PIM135
PIM137
PIM172
PIM147
PIM161
PIM163
PIM164
PIM165
PIM171
PIM166
PIM177
PIM173
PIM180
PIM181
PIM182
PIM190
PIM231
PIM191
PIM195
PIM201
PIM215
PIM214
PIM224
PIM226
PIM225
PIM228
PIM239
PIM244
PIM245
PIM242
PIM269
PIM291
PIM254
PIM311
PIM263
PIM262
PIM284
PIM321
PIM283
PIM285
PIM300
PIM328
PIM326
PIM332
PIM333
PIM334
PIM339
PIM344
PIM346
PIM350
PIM359
PIM358
PIM371
PIM369
PIM373
PIM384
PIM411
PIM397
PIM387
PIM402
PIM407
PIM388
PIM393
PIM404
Chromosome
Copy Number
0 1 2 3 4

### Figure S4

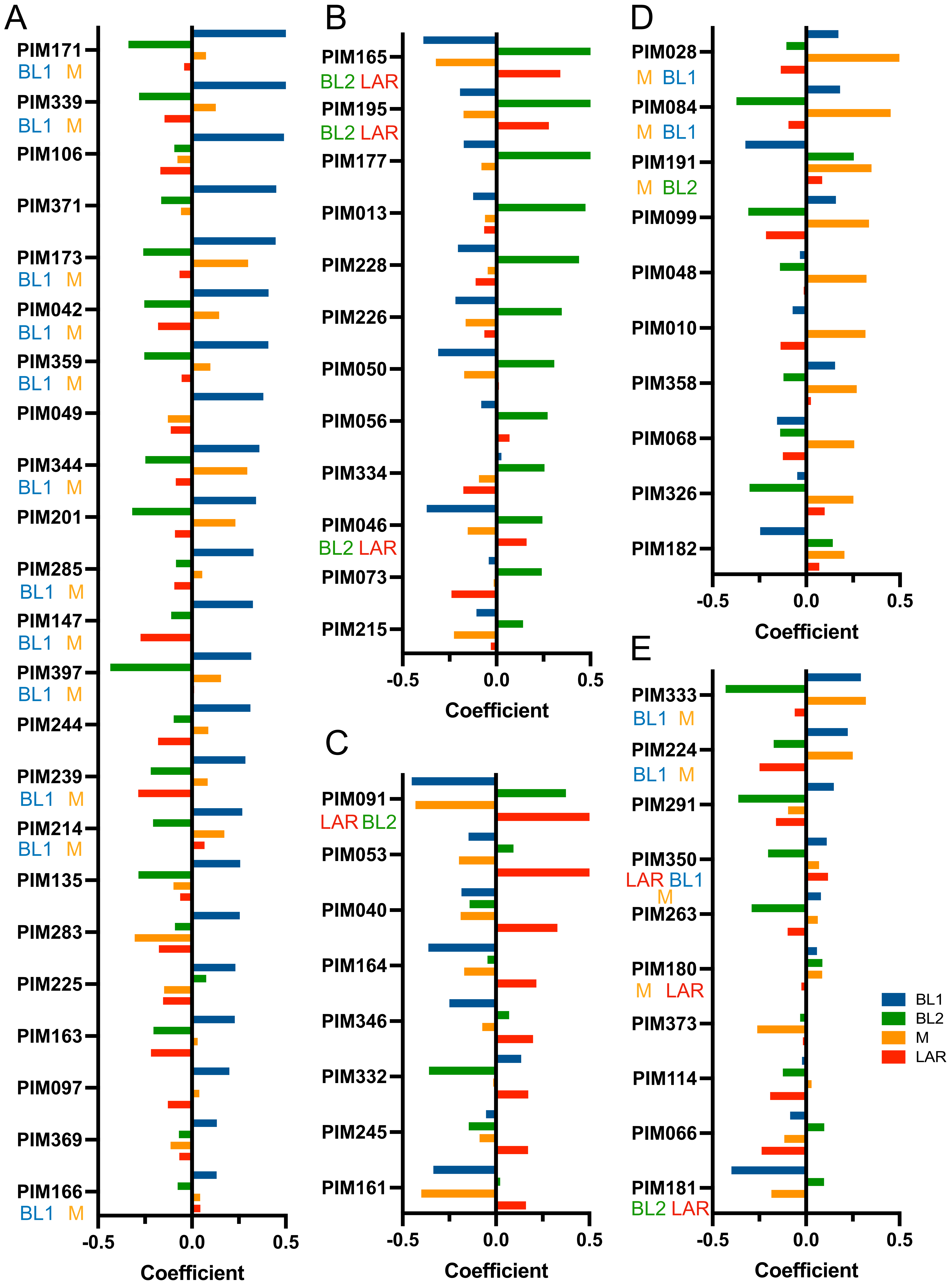

A
PIM171
BL1 M
PIM339
BL1 M
PIM106
PIM371
PIM173
BL1 M
PIM042
BL1 M
PIM359
BL1 M
PIM049
PIM344
BL1 M
PIM201
PIM285
BL1 M
PIM147
BL1 M
PIM397
BL1 M
PIM244
PIM239
BL1 M
PIM214
BL1 M
PIM135
PIM283
PIM225
PIM163
PIM097
PIM369
PIM166
BL1 M
-0.5 0.0 0.5
Coefficient
B
PIM165
BL2 LAR
PIM195
BL2 LAR
PIM177
PIM013
PIM228
PIM226
PIM050
PIM056
PIM334
PIM046
BL2 LAR
PIM073
PIM215
-0.5 0.0 0.5
Coefficient
C
PIM091
LAR BL2
PIM053
PIM040
PIM164
PIM346
PIM332
PIM245
PIM161
-0.5 0.0 0.5
Coefficient
D
PIM028
M BL1
PIM084
M BL1
PIM191
M BL2
PIM099
PIM048
PIM010
PIM358
PIM068
PIM326
PIM182
-0.5 0.0 0.5
Coefficient
E
PIM333
BL1 M
PIM224
BL1 M
PIM291
PIM350
LAR BL1
M
PIM263
PIM180
M LAR
PIM373
PIM114
PIM066
PIM181
BL2 LAR
-0.5 0.0 0.5
Coefficient
BL1
BL2
M
LAR

### Figure S5

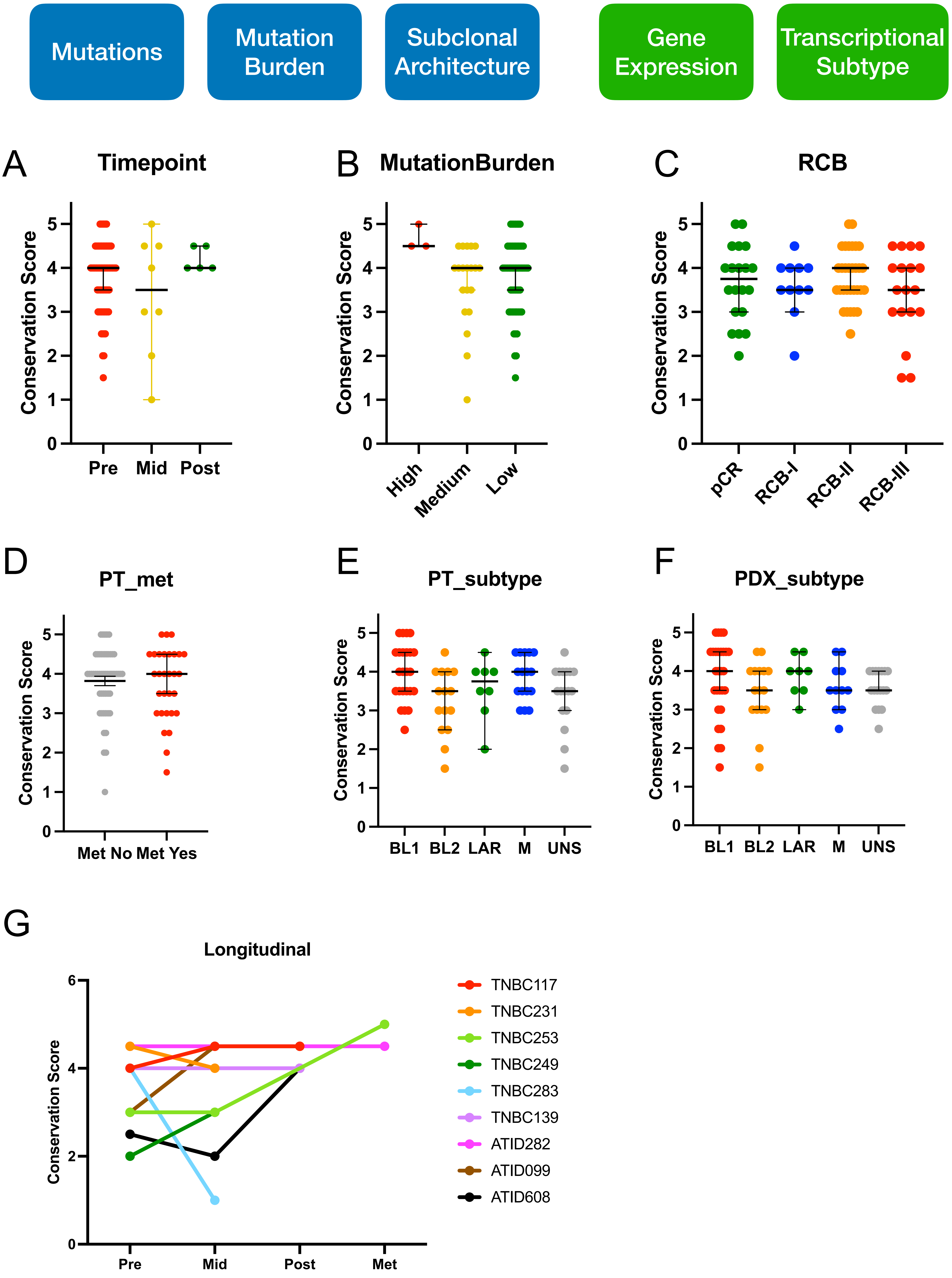

Mutations
Mutation Burden
Subclonal Architecture
Gene Expression
Transcriptional Subtype
A
Timepoint
Conservation Score
Pre
Mid
Post
B
MutationBurden
Conservation Score
High
Medium
Low
C
RCB
Conservation Score
pCR
RCB-I
RCB-II
RCB-III
D
PT_met
Conservation Score
Met No
Met Yes
E
PT_subtype
Conservation Score
BL1
BL2
LAR
M
UNS
F
PDX_subtype
Conservation Score
BL1
BL2
LAR
M
UNS
G
Longitudinal
Conservation Score
Pre
Mid
Post
Met
TNBC117
TNBC231
TNBC253
TNBC249
TNBC283
TNBC139
ATID282
ATID099
ATID608

### Figure S6

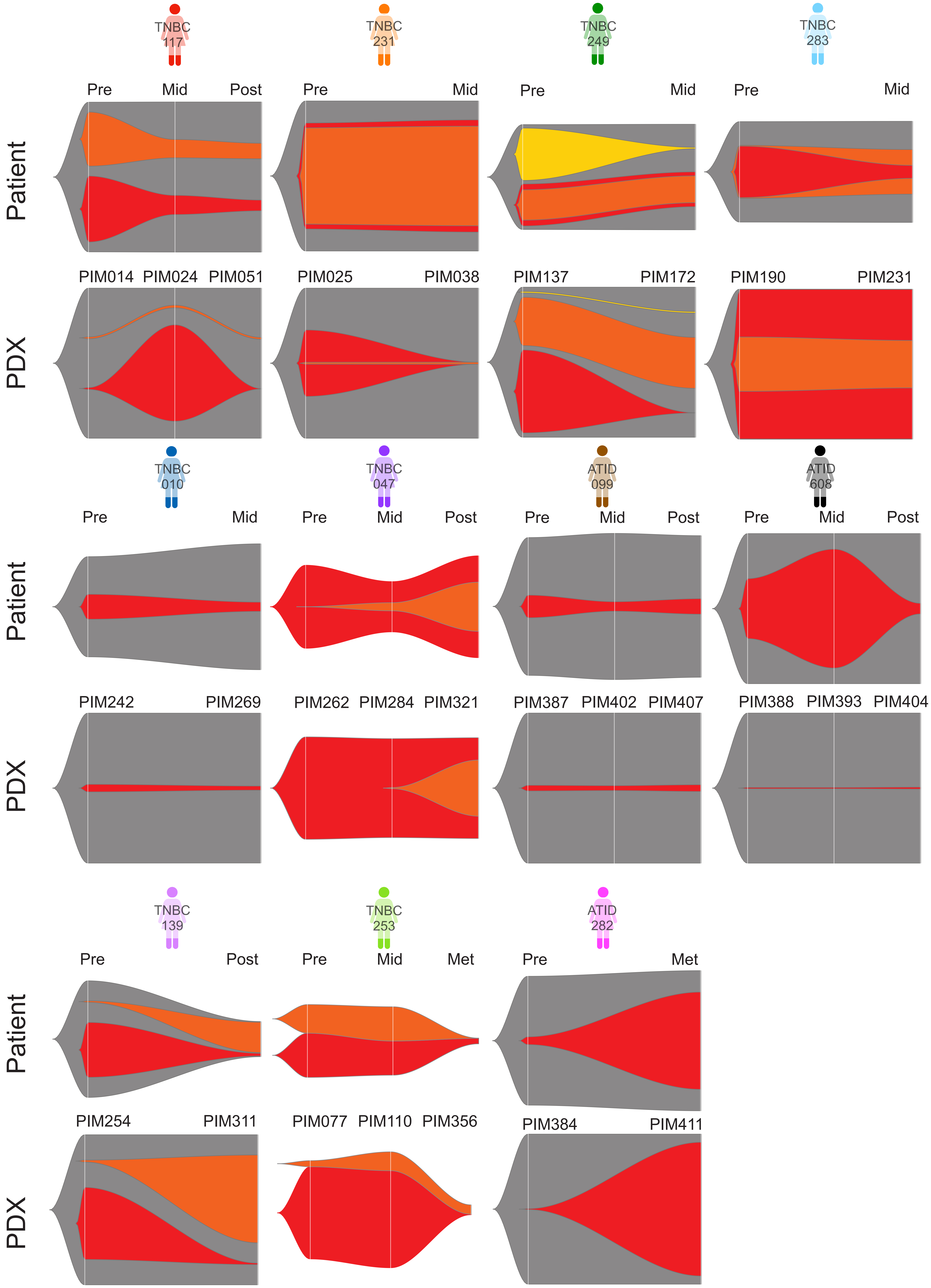

TNBC 117
TNBC 231
TNBC 249
TNBC 283
Pre
Mid
Post
Patient
PIM014
PIM024
PIM051
PIM025
PIM038
PIM137
PIM172
PIM190
PIM231
PDX
TNBC 010
TNBC 047
ATID 099
ATID 608
PIM242
PIM269
PIM262
PIM284
PIM321
PIM387
PIM402
PIM407
PIM388
PIM393
PIM404
TNBC 139
TNBC 253
ATID 282
Met
PIM254
PIM311
PIM077
PIM110
PIM356
PIM384
PIM411

**Figure S7**

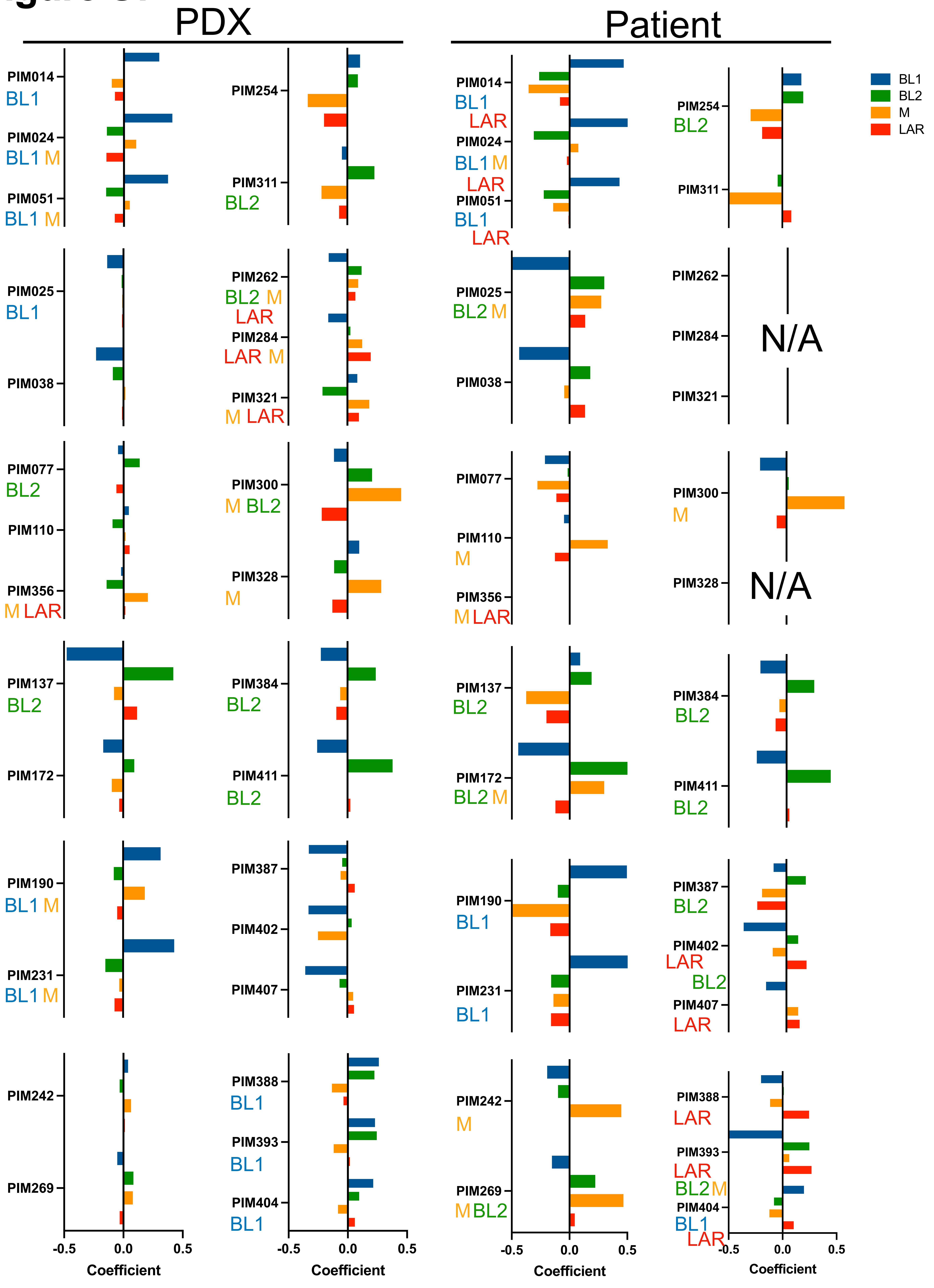
PDX
Patient
BL1
BL2
M
LAR
PIM014
BL1
PIM024
BL1 M
PIM051
BL1 M
PIM254
PIM311
BL2
PIM014
BL1
LAR
PIM024
BL1 M
LAR
PIM051
BL1
LAR
PIM254
BL2
PIM311
PIM025
BL1
PIM038
PIM262
BL2 M
LAR
PIM284
LAR M
PIM321
M LAR
PIM025
BL2 M
PIM038
PIM262
PIM284
PIM321
N/A
PIM077
BL2
PIM110
PIM356
M LAR
PIM300
M BL2
PIM328
M
PIM077
PIM110
M
PIM356
M LAR
PIM300
M
PIM328
N/A
PIM137
BL2
PIM172
PIM384
BL2
PIM411
BL2
PIM137
BL2
PIM172
BL2 M
PIM384
BL2
PIM411
BL2
PIM190
BL1 M
PIM231
BL1 M
PIM387
PIM402
PIM407
PIM190
BL1
PIM231
BL1
PIM387
BL2
PIM402
LAR
BL2
PIM407
LAR
PIM242
PIM269
PIM388
BL1
PIM393
BL1
PIM404
BL1
PIM242
M
PIM269
M BL2
PIM388
LAR
PIM393
LAR
BL2 M
PIM404
BL1
LAR
-0.5
0.0
0.5
Coefficient

### Figure S8

A

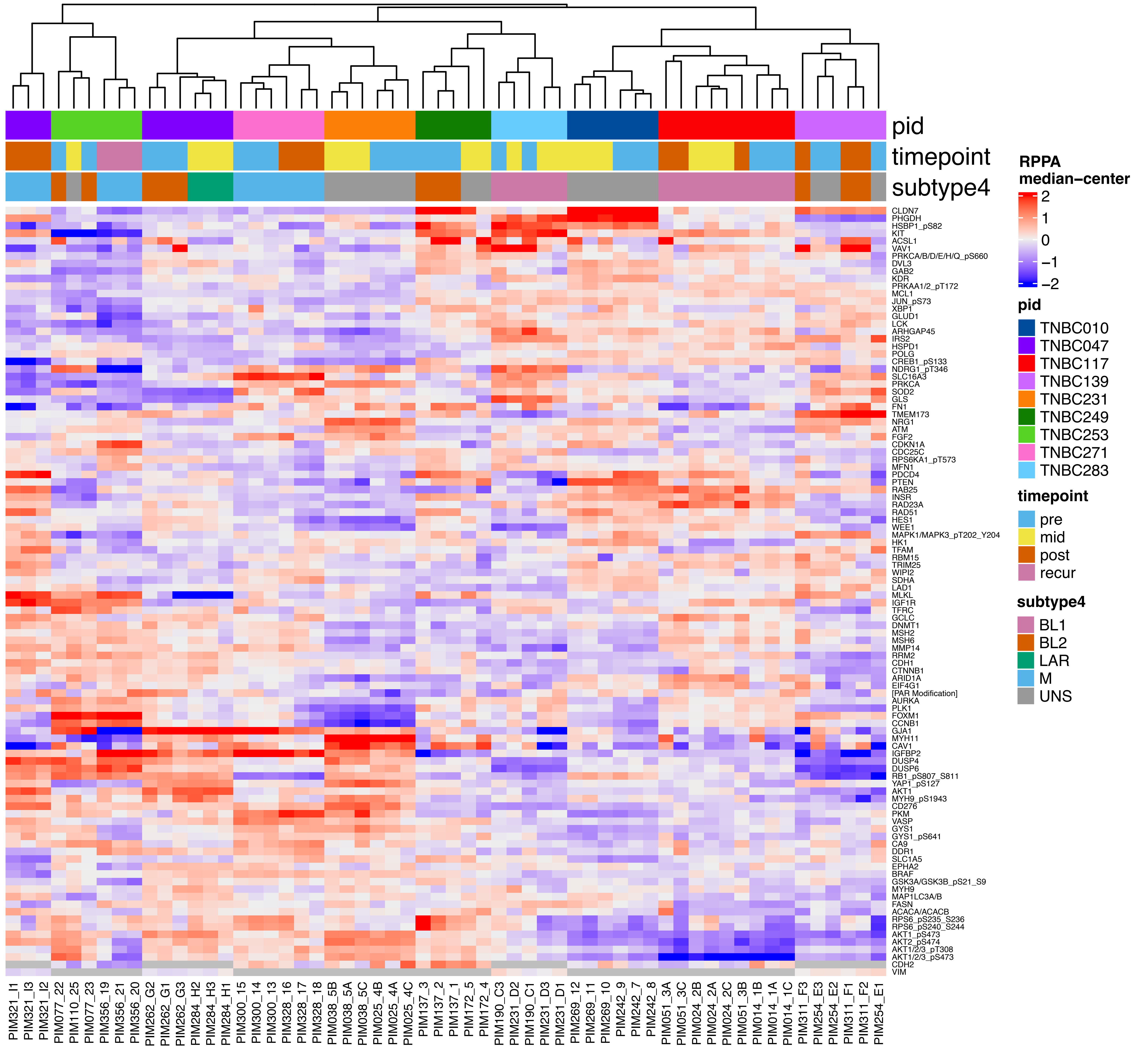

pid
timepoint
subtype4
RPPA median−center
pid
TNBC010
TNBC047
TNBC117
TNBC139
TNBC231
TNBC249
TNBC253
TNBC271
TNBC283
timepoint
pre
mid
post
recur
subtype4
BL1
BL2
LAR
M
UNS

B

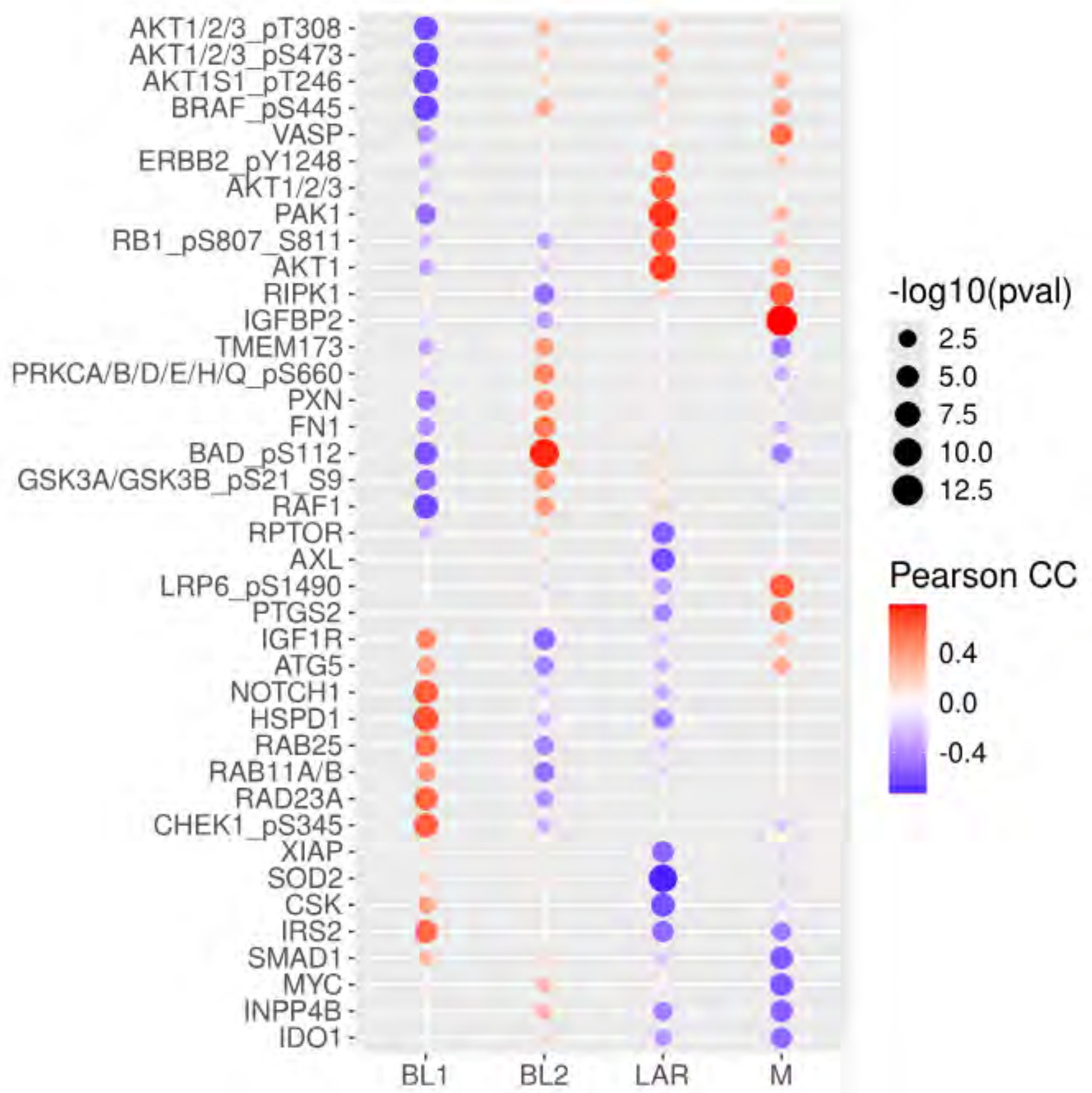

AKT1/2/3_pT308
AKT1/2/3_pS473
AKT1S1_pT246
BRAF_pS445
VASP
ERBB2_pY1248
AKT1/2/3
PAK1
RB1_pS807_S811
AKT1
RIPK1
IGFBP2
TMEM173
PRKCA/B/D/E/H/Q_pS660
PXN
FN1
BAD_pS112
GSK3A/GSK3B_pS21_S9
RAF1
RPTOR
AXL
LRP6_pS1490
PTGS2
IGF1R
ATG5
NOTCH1
HSPD1
RAB25
RAB11A/B
RAD23A
CHEK1_pS345
XIAP
SOD2
CSK
IRS2
SMAD1
MYC
INPP4B
IDO1
BL1
BL2
LAR
M
-log10(pval)
2.5
5.0
7.5
10.0
12.5
Pearson CC
0.4
0.0
-0.4

**Table S2: Summary of available genomic and transcriptomic data**

| | PDX samples (cases) | Patient samples (cases) |
|---|---|---|
| **RNASeq** | 92 (75) | 84 (70) |
| **DNASeq** | 92 (75) | 87 (71) |
| **Common cases** | 92 (75) | 82 (68) |
| **Timepoint** | | |
| **Pre-treatment** | 71 | 67/68* |
| **Mid-treatment** | 11 | 9/11* |
| **Post-treatment** | 8 | 6 |
| **Recurrence** | 2 | 2 |
| **Longitudinal Sets** | 12 | 10/11* |

**Table S3: Summary of available data for longitudinal PDX sets**

| Public ID | PDX | Timepoint | Treatment | RCB | Pt WES | Pt RNA | PDX WES | PDX RNA | PDX RPPA | PDX AC | Drug Screen |
|---|---|---|---|---|---|---|---|---|---|---|---|
| TNBC117 | PIM014 | Pre | None | RCB-III | X | X | X | X | X | | X |
| | PIM024 | Mid | AC | RCB-III | X | X | X | X | X | | X |
| | PIM051 | Post | AA | RCB-III | X | X | X | X | X | | X |
| TNBC231 | PIM025 | Pre | None | RCB-II | X | X | X | X | X | X | X |
| | PIM038 | Mid | AC | RCB-II | X | X | X | X | X | X | X |
| TNBC253 | PIM077 | Pre | None | RCB-II | X | X | X | X | X | | X |
| | PIM110 | Mid | AC | RCB-II | X | X | X | X | X | | |
| | PIM356 | Met | PCT | RCB-II | X | X | X | X | X | | |
| TNBC249 | PIM137 | Pre | None | pCR | X | X | X | X | X | X | X |
| | PIM172 | Mid | AC | pCR | X | X | X | X | X | X | X |
| TNBC283 | PIM190 | Pre | None | ND | X | X | X | X | X | X | X |
| | PIM231 | Mid | AC | ND | X | X | X | X | X | X | X |
| TNBC010 | PIM242 | Pre | None | RCB-II | X | X | X | X | X | X | X |
| | PIM269 | Mid | AC | RCB-II | X | X | X | X | X | X | X |
| TNBC139 | PIM254 | Pre | None | RCB-II | X | X | X | X | X | X | X |
| | PIM311 | Post | AC, P | RCB-II | X | X | X | X | X | X | X |
| TNBC047 | PIM262 | Pre | None | RCB-II | X | | X | X | X | X | X |
| | PIM284 | Mid | AC | RCB-II | X | | X | X | X | X | X |
| | PIM321 | Post | AA | RCB-II | X | | X | X | X | X | X |
| TNBC271 | PIM300 | Pre | None | RCB-II | X | X | X | X | X | X | X |
| | PIM328 | Post | AA | RCB-II | | | X | X | X | X | X |
| ATID282 | PIM384 | Pre | None | RCB-II | X | X | X | X | | | |
| | PIM411 | Met | AC, P | RCB-II | X | X | X | X | | | |
| ATID099 | PIM387 | Pre | None | RCB-III | X | X | X | X | | | X |
| | PIM402 | Mid | AC | RCB-III | X | X | X | X | | | X |
| | PIM407 | Post | PE | RCB-III | X | X | X | X | | | X |
| ATID608 | PIM388 | Pre | None | RCB-II | X | X | X | X | | X | X |
| | PIM393 | Mid | AC | RCB-II | X | X | X | X | | X | X |
| | PIM404 | Post | PCT | RCB-II | X | X | X | X | | X | X |

Abbreviations: AC – doxorubicin and cyclophosphamide, AA – nab-paclitaxel and atezolizumab, P – paclitaxel, PCT – paclitaxel, carboplatin, panitumumab, PE – paclitaxel, enzalutamide, RCB – residual cancer burden, pCR – pathological complete response, WES – whole exome sequencing, RPPA – reverse phase protein array
